# Cancer subclone detection based on DNA copy number in single cell and spatial omic sequencing data

**DOI:** 10.1101/2022.07.05.498882

**Authors:** Chi-Yun Wu, Anuja Sathe, Jiazhen Rong, Paul R. Hess, Billy T. Lau, Susan M. Grimes, Hanlee P. Ji, Nancy R. Zhang

## Abstract

In cancer, somatic mutations such as copy number alterations (CNAs) accumulate during disease progression and lead to functional intra-tumor heterogeneity that can influence the efficacy of cancer therapy. Therefore, studying the functional characteristics and spatial distribution of genetically distinct subclones is crucial to the understanding of tumor evolution and the design of cancer treatment. Here, we present Clonalscope, a method for subclone detection using copy number profiles that can be applied to spatial transcriptomics (ST) data and data from single-cell sequencing platforms such as scRNA-seq and scATAC-seq. Clonalscope implements a nested Chinese restaurant process to identify *de novo* subclones within one or multiple samples from the same patient. Clonalscope incorporates prior information from paired whole-genome or whole-exome sequencing (WGS/WES) data to achieve more reliable subclone detection and malignant cell labeling. On scRNA-seq and scATAC-seq data from four gastrointestinal tumor samples, Clonalscope successfully labeled malignant cells and identified genetically different subclones, which were validated in detail using matched scDNA-seq data. On ST data from a squamous cell carcinoma and two invasive ductal carcinoma samples, Clonalscope successfully labelled malignant spots, traced subclones between associated datasets, and identified spatially segregated subclones expressing genes associated with drug resistance and survival.

## Introduction

Cancer is a disease of intra-tissue clonal evolution, where DNA mutations combine with epigenetic alterations to confer selective advantages on neoplastic cells, allowing them to proliferate, invade, metastasize, and resist therapeutic treatment ^1–2^. Detection and characterization of genetically different subclones is crucial for the study of cancer and for developing effective treatments and prognostic strategies. Copy number alteration (CNA), one type of genomic variation, is a prevalent manifestation of genome instability and a key driver of tumor evolution ^3–5^ Subclones distinguished by CNA have been successfully profiled using bulk and single-cell DNA sequencing data ^6–14^ CNA-based subclone detection in other types of single-cell omic data, such as single-cell RNA sequencing (scRNA-seq) ^15–19^ and single-cell transposase-accessible chromatin-sequencing (scATAC-seq) ^20, 21^ further allows the joint analysis of genetic mutations with gene expression and chromatin state. Recently, the maturation of spatial transcriptomic (ST) technologies ^22, 23^ is heralding new opportunities for the study of spatial subclonal dynamics through the mapping of DNA alterations of cells in situ ^24, 25^.

There is a need for reliable methods for CNA estimation and subclone detection for scRNA-seq, scATAC-seq and ST data. Since these methods do not profile DNA, read coverage along the chromosome is, at best, a noisy and indirect proxy of the underlying DNA copy number. Existing methods for scRNA-seq data rely on smoothing across adjacent genes, e.g. inferCNV ^16^ and CopyKAT ^15^. Similar strategies were also used in CNA analysis for scATAC-seq data ^21^. Other methods such as HoneyBADGER ^18^ and CaSpER ^17^ consider allelic expression in their copy number detection model, but the sparsity of allele-specific reads limits the improvement achieved by these approaches. For spatial transcriptomics data, there is yet no vetted published method for CNA-based subclone detection. STARCH, developed for CNA detection for ST data, leverages the signals from nearby spots to overcome sparsity ^25^. Since scRNA-seq and ST both measure gene expression, some emerging studies apply inferCNV, developed for scRNA-seq data, to analyze CNA in tumor ST data ^24^.

The above methods partition cells/spots into clusters with distinct transcriptomic/epigenetic profiles, but to what degree do the clusters represent subclones with distinct DNA-level copy number profiles? Since broad megabase-level shifts in accessibility/transcription have been observed between normal cells of different lineages, and may reflect chromatin-level remodeling instead of copy number, RNA-based CNA detection can be prone to false-positives. In contrast, CONICS ^19^ profiles CNAs in scRNA-seq data with segmentation obtained from matched bulk DNA-seq data to reduce false signals, but it was mainly developed for the full-length scRNA-seq data and does not infer intratumor subclones. Like the above approaches, CONICS relies on hierarchical clustering to cluster cells into subclones. Hierarchical clustering works when there are clear, well-separated clusters in the data, and breaks down when the signal-to-noise ratio is low as in the case of copy number estimates from scRNA-seq, scATAC-seq, and ST data. Hierarchical clustering does not leverage the prior information in matched bulk DNA sequencing data, which is often available, nor does it leverage the specific structure of subclonal copy number changes that could be used to distinguish signal from noise.

We present Clonalscope, a method for CNA-based subclone detection that can be applied to diverse single cell and spatial sequencing data types, such as scRNA-seq, scATAC-seq, and spatial transcriptomics data. Clonalscope is based on nonparametric Bayesian clustering via a novel nested Chinese restaurant process, which exploits the sparse nature of subclonal CNA differences to identify *de novo* subclones with related, but distinct, copy number profiles. Importantly, Clonalscope allows the leveraging of prior information from paired whole-genome or whole-exome sequencing (WGS/WES) data to achieve more reliable subclone detection and malignant cell labeling. When multiple samples are sequenced for a patient, Clonalscope also allows the linking of subclones across samples for lineage tracing.

We first consider CNA estimation, malignant cell labeling and cancer subclone detection in scRNA-seq data, where tissue-matched bulk and single-cell DNA sequencing data are used for benchmarking. We show that CNA estimates derived from scRNA-seq data alone, without leveraging matched bulk DNA-seq data, can be inaccurate, and that the errors propagate downstream to affect such common tasks as malignant cell identification. Leveraging WGS/WES data, Clonalscope obtains precise copy number estimation and significantly improves the accuracy of malignant cell labeling and subclone detection, as compared to existing approaches. Next, we apply Clonalscope to scATAC-seq data from a gastric cancer cell line with matched scDNA-seq data for validation, and show that Clonalscope can identify subclones based on allele-specific copy number profiles. Finally, on ST data from breast and skin cancer, with expert pathology annotation as validation, we show that Clonalscope successfully distinguishes malignant from non-malignant cells and achieves reproducible spatial subclone mapping across adjacent tissue slices.

## Results

### Overview of the Clonalscope model and algorithm (Fig. 1)

Clonalscope starts by segmenting the genome into regions of homogeneous copy number states in matched bulk whole-genome sequencing (WGS) or whole-exome sequencing (WES) data (Step. 1). Next, in single-cell RNA/ATAC sequencing (scRNA-seq/scATAC-seq) data, the relative coverage fold change for each cell *i* in each region *r* is estimated under a Poisson model with the baseline coverage computed from a set of pre-chosen diploid cells, such as immune cells (Step. 2; Methods). This reduces the cell-by-gene matrix to a cell-by-region matrix, which serves as input to the non-parametric Bayesian clustering model for subclone detection and subclonal copy number inference (Step. 3). The non-parametric Bayesian clustering model is initialized to two clusters: a cluster representing normal cells with fold change initialized to 1 across all regions, and a cluster representing the major tumor clone with fold change in each region initialized to its fold change in the matched bulk DNA sequencing (DNA-seq) data. Then, Clonalscope applies a nested Chinese Restaurant Process to iteratively add/select copy number states for each genome segment, seed new subclones, and assign each cell to a subclone. We use the term “nested” since the algorithm involves two Chinese Restaurant Processes: The first Chinese Restaurant Process samples subclone membership and seeds new subclones, while for each new cancer subclone, a second Chinese Restaurant Process samples the copy number of each region, seeding new copy number states as needed. Clonalscope outputs the estimated numbers of subclones, the copy number profiles for the subclones, and the subclone assignment of each cell.

**Figure 1:**
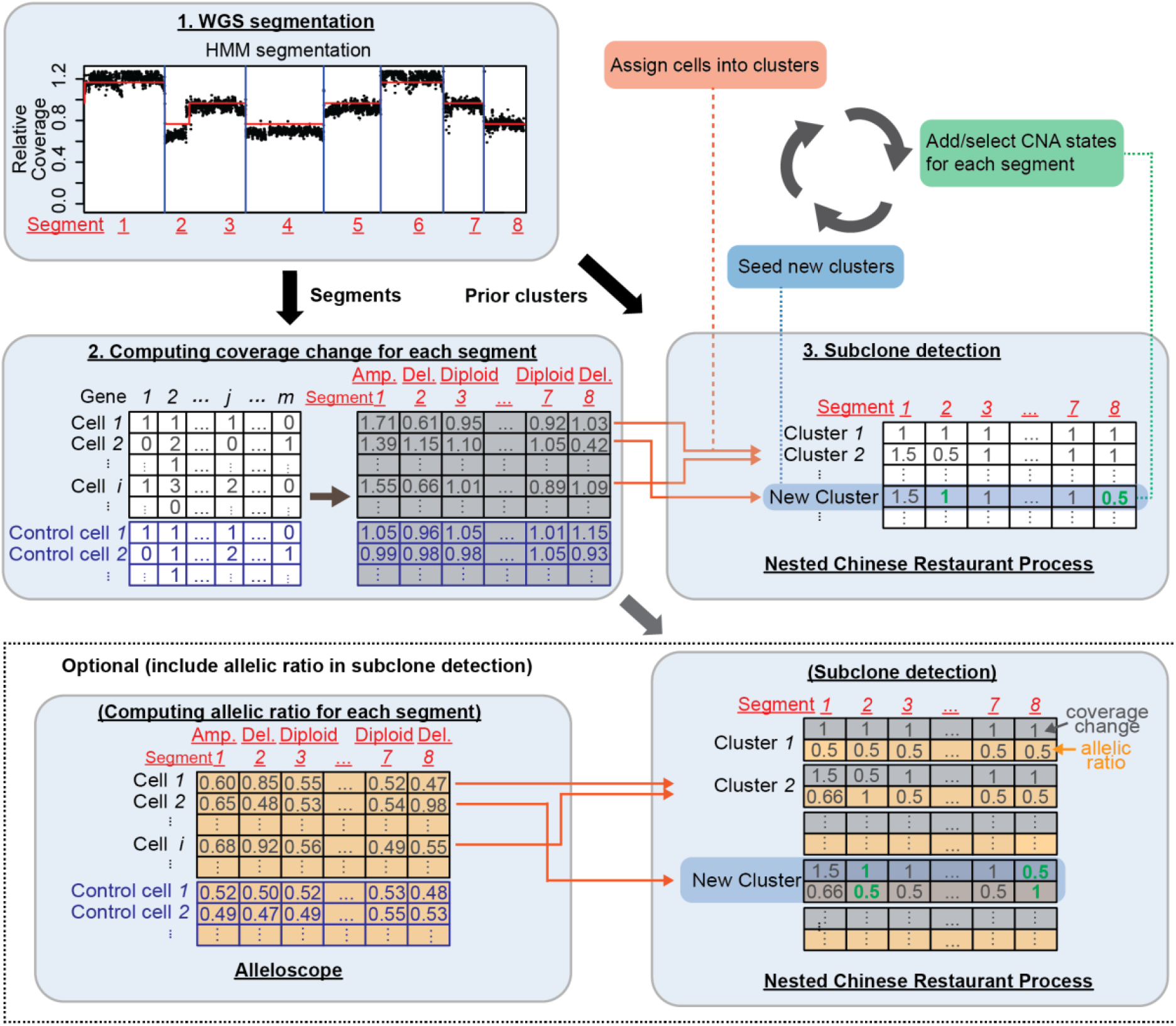
Overview of subclone detection with Clonalscope. Step 1. The algorithm starts from a segmentation of the genome based on matched WGS or WES data. Step 2. For each segment, Clonalscope computes the coverage change of each cell in scRNA-seq data by normalizing to a set of normal cells. This step reduces the raw gene-by-cell matrix for the UMI counts to the segment-by-cell matrix. Step 3. Clonalscope implements a nested Chinese Restaurant Process to iteratively assign each cell into one of the existing clusters, add or select copy number states for each segment (1^st^ Chinese Restaurant Process), and seeds new clusters (2^nd^ Chinese Restaurant Process) (Methods). For ST data, the setting is similar but “cell” is substituted with “spot”. Optionally, Clonalscope can also operate on both coverages and allelic ratios for subclone detection on scATAC-seq data. In addition to the segment-by-cell matrix for fold changes, Alleloscope can be used to retrieve the segment-by-cell matrix for allelic ratios. Clonalscope then detects subclones based on both types of information.

For spatial transcriptomic (ST) data, the same algorithm as described above applies, with “cell” replaced by “spot”. We do not make use of the spatial coordinates in the clustering, and we don’t smooth the cluster memberships across space. This is because we do not want to make a priori assumptions on the degree of intermixing of subclones in space.

Clonalscope can also make use of allele-specific coverage, if such data is available. As above, we start with a segmentation of the genome derived from bulk DNA-seq data, and then use Alleloscope ^9^ to estimate both the relative fold change and allelic ratio for each cell *i.* in each region *r.* The non-parametric Bayesian clustering then considers both relative fold change and allelic ratio in computing the likelihood of a cell given a copy number profile. See Methods for details.

### Subclone detection in scRNA-seq data validated by matched scDNA-seq data

Most existing copy number aberration (CNAs) estimation methods for scRNA-seq data do not allow integration with bulk WGS data, and rely solely on a smoothing/averaging of the RNA expression data for copy number estimation. Then, these methods usually apply hierarchical clustering on the estimated single cell copy number profiles for subclone detection or malignant cell labeling. There can be considerable differences between RNA expression and DNA copy number ^26^, even when the former is smoothed across large contiguous regions ^27^. To examine how well scRNA-seq data can recapitulate the underlying *DNA-level* heterogeneity, we consider scRNA-seq data from three primary gastrointestinal cancer samples (P5847, P5931, and P8823) and one colorectal metastasis (P6198). For P5847 and P5931, scDNA-seq data is available for validation, and for P6198 and P8823, bulk DNA-seq is available for assessment of concordance of mean copy number profiles. We benchmarked Clonalscope against inferCNV and CopyKAT, two commonly used methods. First, we compared the correlations between mean copy number estimates derived from epithelial cells in the scRNA-seq data with copy number estimates derived from the matched DNA-seq data (Fig. 2a). All epithelial cells in the metastasized tumor P6198 are malignant cells. The three primary tumors (P5847, P5931, and P8823) have high purity and thus most of the epithelial cells are expected to be malignant. Therefore, we expect high correlation between the CNA profiles of epithelial cells in scRNA-seq data and that from the matched DNA-seq data. Across the four samples (Supplementary Table 1), correlations of Clonoscope estimates all exceed 0.7, while existing methods which rely solely on scRNA-seq data achieve correlations around 0.5. The detailed copy number estimates are shown in Supplementary Fig. 1. Across the four samples, the lack of concordance between copy number estimates obtained by existing methods and gold standard values derived from DNA-seq demonstrates the difficulty of copy number estimation solely from RNA expression, and motivates the integration with matched bulk DNA-seq for copy number-based subclone detection in scRNA-seq data.

**Figure 2:**
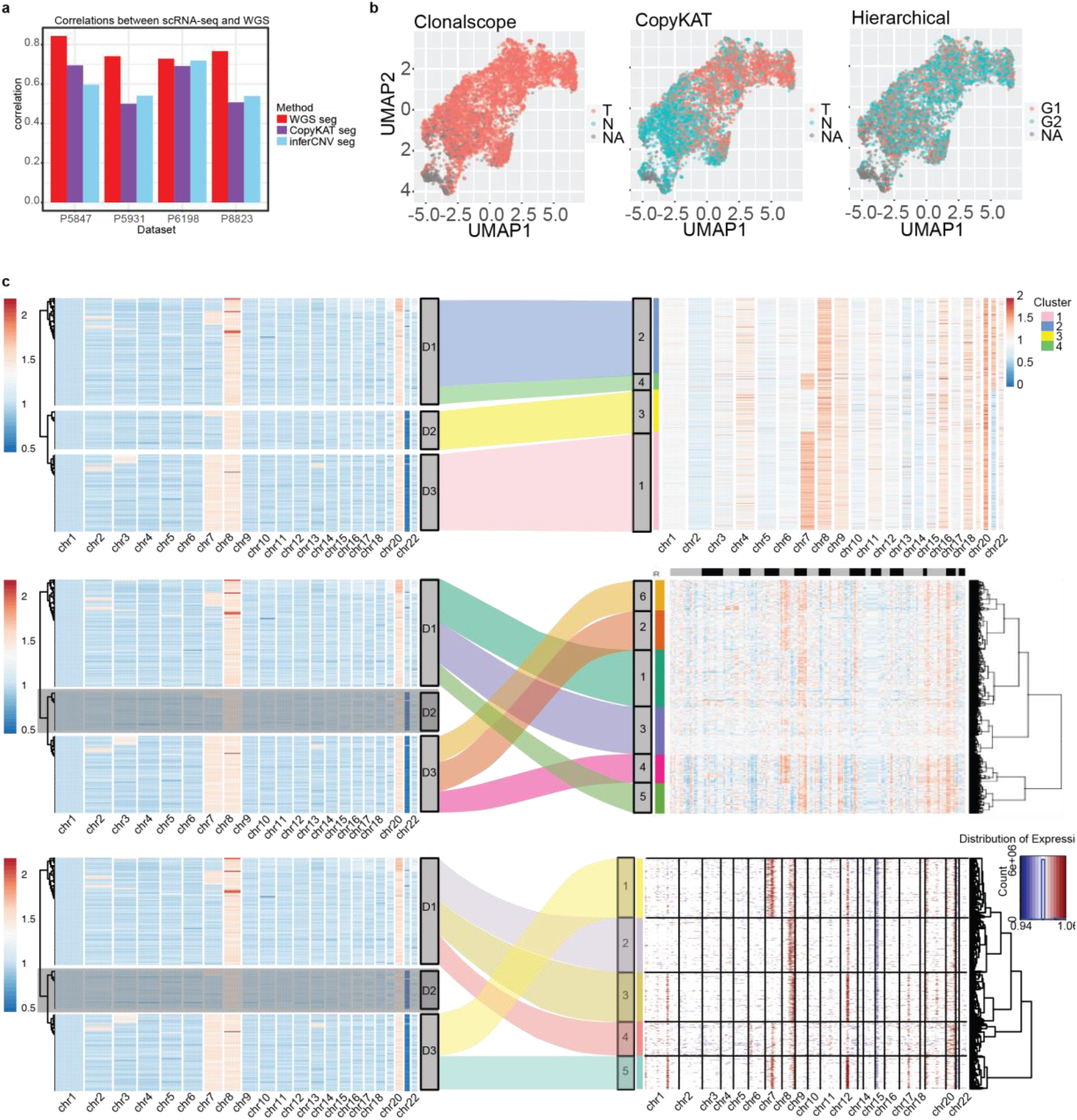
Subclone detection and malignant cell labeling for scRNA-seq data validated by matched scDNA-seq data. a. Correlations between the copy number profiles from matched DNA-seq data and those from different segmentation methods on four gastrointestinal tumor samples. b. Malignant cell labeling from Clonalscope (left), CopyKAT (middle), and Hierarchical clustering with 2 major groups (right) on the P6198 scRNA-seq data. T and N represent the estimated tumor and normal cells respectively. G1 and G2 stand for the first and second major groups from hierarchical clustering. c. Mapping the detected subclones from Clonalscope (top), CopyKAT (middle), and inferCNV(bottom) in scRNA-seq data (right) to those observed in the matched scDNA-seq data (left) for the P5931 malignant cells (Methods).

In addition to providing the genome segmentation which serves as the starting point of Clonoscope’s single-cell subclone detection, the copy number profile from tissue-matched bulk DNA-seq data is also informative for the assignment of scRNA-seq cells to malignant versus non-malignant status, a common analysis task in single cell cancer sequencing. Existing methods use hierarchical clustering for this task, without utilizing the tissue-matched bulk DNA-seq data. Consider P6198, the metastasized tumor from above where all epithelial cells are known to be of malignant status and thus can be used to benchmark copy number-based malignant cell identification accuracy. The results of copy number-based malignant cell labeling by Clonalscope (left), CopyKAT (middle) and hierarchical clustering (right) are shown in Fig. 2 (inferCNV does not perform malignant cell assignment). By utilizing the CNA profile from matched bulk DNA-seq data, Clonalscope’s accuracy in malignant cell labeling reaches ~0.9986 accuracy. In comparison, CopyKAT’s malignant cell labeling, which does not utilize matched bulk DNA-seq data, has accuracy of ~0.4529 on this sample (Supplementary Fig. 2). While we recognize that tissue-matched bulk DNA-seq data is not always available, this example shows that when it is available, it can be harnessed by Clonalscope to substantially boost accuracy. The accuracy boost is derived not just from the DNA-based segmentation, but also from Clonalscope’s clustering model which utilize the bulk DNA profile as prior. This can be seen from the fact that even when the segmentation from matched bulk DNA-seq data is used to delineate the CNA regions, hierarchical clustering is still not able to reliably label malignant cells (Supplementary Fig. 3).

Next, we compared the accuracy of subclone identification by Clonalscope, CopyKAT and inferCNV for scRNA-seq data. Existing methods for copy number-based subclone detection for scRNA-seq data have not explicitly evaluated the accuracy of intratumor subclones using scDNA-seq gold standard. Instead, benchmarking was only done at the bulk level. The comparison here was performed on the P5931 primary gastric tumor sample since it has a clear subclonal structure from scDNA-seq data (Fig. 2c left and ^9^), and furthermore, it has matched normal scRNA-seq data generated for benchmarking the performance of subclone identification (Supplementary Fig. 4). Focusing on the malignant cells in the P5931 scRNA-seq dataset, the subclones identified from Clonalscope (top), CopyKAT (middle), and inferCNV (bottom) were compared to the three main subclones (D1 to D3) observed in matched scDNA-seq data (Fig. 2c-e; Methods). Proportions of each subclone detected by each method are shown in Supplementary Fig. 5. Clonalscope successfully identified all three major subclones with proportions comparable to those in scDNA-seq data (Supplementary Fig. 5). Comparatively, subclone D3 was apparently missed in the results of CopyKAT and inferCNV (Fig. 2d, e). Also, compared to existing methods, Clonalscope does not require stringent filters and thus retains more low coverage cells in the analysis (Supplementary Fig. 5). Thus, with validation using scDNA-seq data, we have shown that Clonalscope is able to improve the accuracy of malignant cell labeling and intratumor subclones detection. The improvements are achieved by harnessing tissue-matched bulk DNA-seq data and by a nonparametric Bayesian model tailored for clustering of clonal copy number profiles.

### Subclone detection based on coverage and allelic ratio

Clonalscope can also utilize allele-specific information for subclone detection, when enough allele-specific reads can be extracted from single-cell sequencing data. We demonstrate this application on scATAC-seq data for SNU601, a gastric cancer cell line with multiple subclones delineated by scDNA-seq data. Some subclones in SNU601 are differentiated mostly by their allele-specific copy number changes ^9^. In our previous study ^9^, we applied a supervised classification approach to assign each cell into one of the six major subclones detected in the matched scDNA-seq data. If we were, instead, to perform unsupervised hierarchical clustering on the scATAC-seq data, we would not be able to recapitulate the subclones due to the low signal-to-noise ratio (Supplementary Fig. 6). The results for Clonoscope are shown in Fig. 3. Based on the segmentation of the pseudo-bulk (Fig. 3a), cells in the SNU601 scDNA-seq data can be clustered into six major subclones with 10 marker regions shown in Fig. 3b, which can be viewed as the ground truth for evaluating the subclone detection results in the much shallower scATAC-seq data (Fig. 3c).

**Figure 3:**
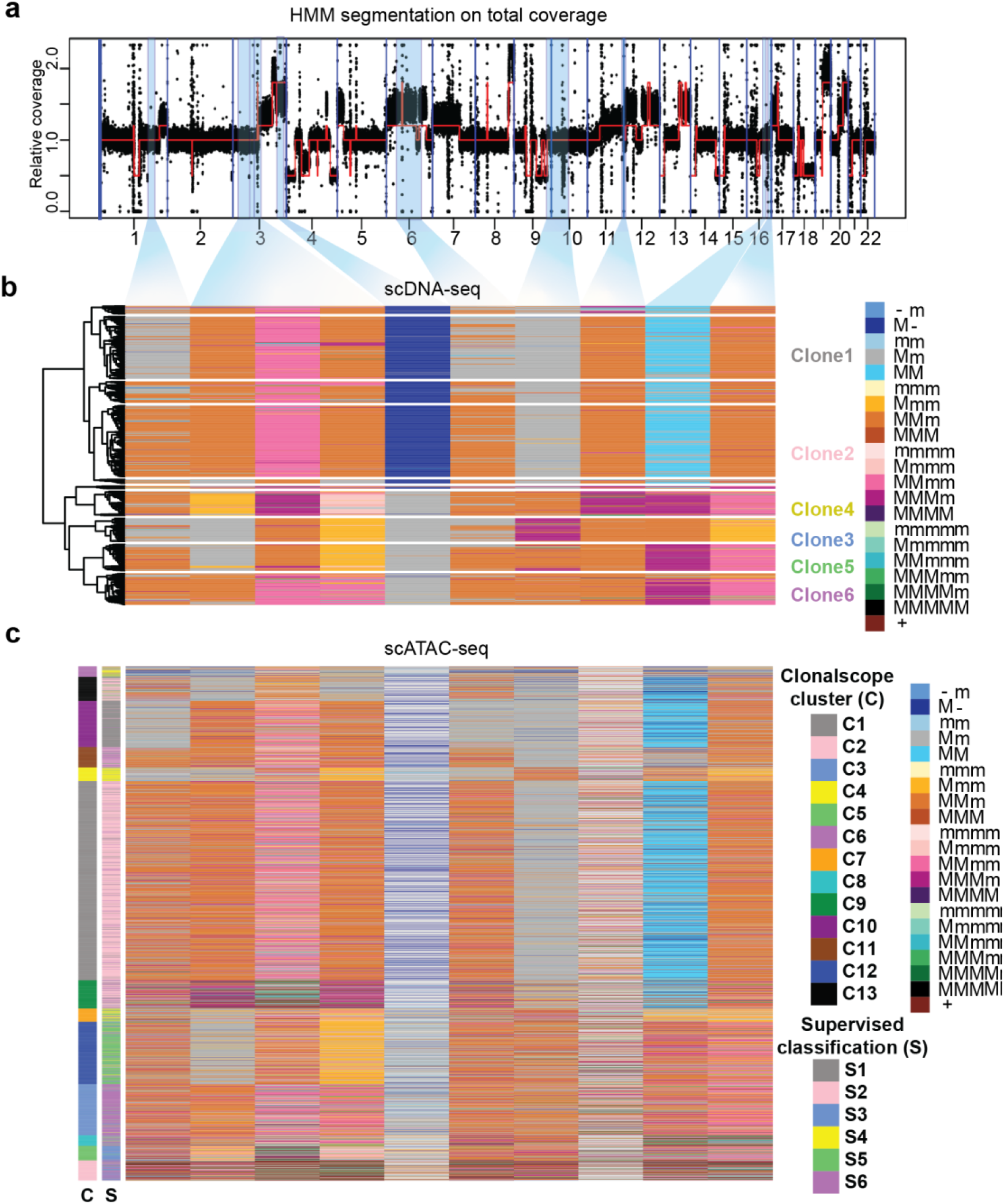
Subclone detection based on coverage and allelic ratio in the scATAC-seq data from the SNU601 gastric cancer cell line. a. Segmentation plot of the pseudo-bulk scDNA-seq dataset from the SNU601 sample. b. The six major subclones detected on the matched scDNA-seq data from hierarchical clustering with ten regions shown in plot c. The result of subclone detection from Clonalscope on the scATAC dataset. The clusters from Clonalscope (C) and supervised classification (S) are shown parallelly in the left. In the color legend, M and m represent the major haplotype and minor haplotype, respectively.

We applied Clonalscope to the scATAC-seq data in the 10 regions in Fig. 3a, and identified 13 subclones in an unsupervised manner (Fig. 3c). The leftmost column shows the subclone identity of each cell assigned by Clonalscope (C). For comparison, we also conducted supervised classification—simply assigning each cell in scATAC-seq to one of the six subclones identified by scDNA-seq, utilizing the subclonal mean copy number profiles from scDNA-seq data. The clusters from supervised classification (S) are also shown in Fig. 3c. A detailed comparison of this supervised subclone assignment versus Clonalscope unsupervised clustering are shown in the contingency table in Supplementary Fig. 7. We see that the six major subclones (right) all have counterparts in Clonalscope’s clusters that exhibited high enrichment. Specifically, the cells assigned to S1 to S6 are mainly enriched in C10, C1, C5, C4, C12, and C3 respectively. The consensus plot also shows comparable allele-specific copy number profiles between the mapped clones from Clonalscope and those derived from the scDNA-seq data (Supplementary Fig. 8). Additionally, we computed the UMAP coordinates for cells in the scATAC-seq data based on their peak profiles and colored the cells based on their clonal assignment from Clonalscope and from supervised classification (Supplementary Fig. 9; ^9^). Clusters from the two methods show similar local enrichment patterns on the UMAP plots. The example here indicates that Clonalscope can detect subclones not only based on total coverage but also based on integrated modeling of total coverage and allele-specific copy number profiles.

### Malignant spot labeling and subclone tracing on spatial transcriptomic data

We next demonstrate the application of Clonalscope on ST data. First, consider an ST data set consisting of two adjacent slices from a squamous cell carcinoma (SCC) sample, with matched bulk WES data. Leveraging the segmentation and bulk copy number profile from the matched WES data (Fig. 4a), Clonalscope identified malignant cells in both slices, which are spatially concentrated despite the fact that Clonalscope did not make use of spatial coordinates in subclone detection (Fig. 4b). Clonalscope’s malignant spot labeling are highly concordant with expert pathology annotation (Fig. 4c). Further, across the two spatially adjacent slices, Clonalscope’s copy number estimates, for the spots labeled as malignant, are highly correlated (correlation ~0.84 Fig. 4d,e). Since the two spatially adjacent slices serve as approximate technical replicates of each other, this analysis demonstrates the accuracy and reproducibility of Clonoscope in malignant cell labelling for spatial transcriptomic data. Detailed heatmaps are shown in Supplementary Fig. 10.

**Figure 4:**
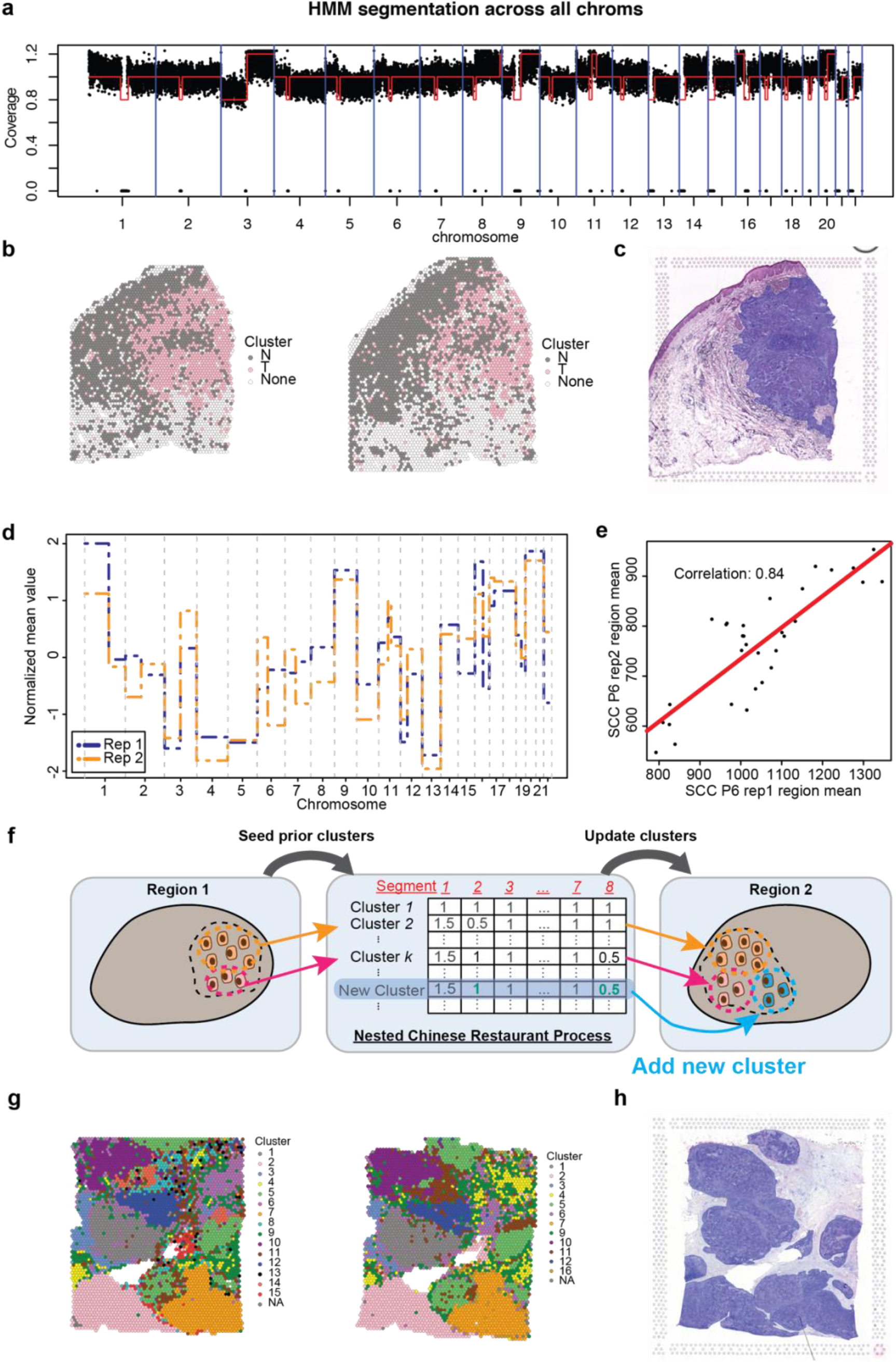
Malignant spot labeling and subclone tracing on spatial transcriptomics (ST) data. a. Segmentation plot of the whole-exome sequencing (WES) data from the squamous cell carcinoma (SCC) Patient 6 (P6) sample. b. Malignant cell labeling by Clonalscope on the ST datasets from SCC P6 replicate 1 (left) and replicate 2 (right). c. The H&E image of the SCC P6 replicate 1 sample with pathology annotation. The regions containing malignant cells are annotated with blue overlays. d. Normalized genome-wide copy number profiles between the identified malignant cells in the two replicates. e. The scatter plot of the mean read counts for each region across the identified malignant cells for SCC P6 replicate 1 against those for replicate 2. f. Illustration of the scheme for subclone detection using Clonalscope. Clonalscope can leverage the prior subclones from the first dataset to detect subclones in another related dataset. g. Subclones detected from Clonalscope in the ST datasets from section 2 (left) and section 1 (right) of an invasive ductal carcinoma (IDC) sample. h. The H&E image of the IDC section 1 sample with pathology annotation. The regions containing malignant cells are annotated with blue overlays.

Currently, there is no method for reliable subclone tracing between samples collected at multiple timepoints, anatomic positions, or spatial slices for the same patient, which is common in today’s study designs. Since Clonalscope allows the flexible integration of prior information in subclone detection, it can be applied to link subclones between samples for the same patient (Fig. 4f). Subclone tracing by Clonalscope is demonstrated on an invasive ductal carcinoma (IDC) data set with samples from two tissue sections (https://www.10xgenomics.com/resources/datasets/human-breast-cancer-block-a-section-1-1-standard-1-0-0). Since there are no bulk DNA sequencing data available for these samples, we simply segmented the genome by chromosome arm. First, we applied Clonalscope on the ST data from one section, yielding a total of 15 clusters (Fig. 4g; left). As before, even though the clustering algorithm does not conduct any smoothing across adjacent spots, the detected subclones are mostly spatially contiguous, which is evidence that the subclones are truly spatially localized. Next, these clusters were used as priors for the subclone detection for the second tissue section, while allowing new clusters to be seeded and prior clusters to be modified/removed (Fig. 4f). In the second tissue section, 11 out of the 15 clusters detected by Clonoscope were preserved with one novel cluster detected (Fig. 4g; right). Comparing the spatial distributions of the detected clusters in the two adjacent sections, we see that the sites of the corresponding clusters match well between the two slides. In both tissue sections, the shapes and positions of subclones delineated by Clonalscope are concordant with the outlines of the tumor in the pathology annotation (Fig. 4h).

### Clonalscope identified spatially segregated subclones expressing genes associated with drug resistance and poor survival

As a second demonstration of the application of Clonalscope to subclone detection in ST data, we analyzed VISIUM 10x data from an invasive ductal carcinoma (IDC sample) (https://www.10xgenomics.com/resources/datasets/human-breast-cancer-ductal-carcinoma-in-situ-invasive-carcinoma-ffpe-1-standard-1-3-0). Since there are no matched bulk WGS data for this sample, copy numbers were estimated at the chromosome arm level. In this sample, Clonalscope identified nine clusters based on the copy number profiles (Supplementary Fig. 11). The nine clusters are locally enriched in the tissue (Fig. 5a) although no spatial information was used in the estimation. Among the clusters, cluster 2 is the major clone; cluster 3 and cluster 4 are the minor subclones spatially segregated in the tissue with expert pathology annotation shown in Supplementary Fig. 12. Most spots containing malignant cells are contained in cluster 2. Comparatively, spots in cluster 3 are only limited to the connected region labeled in blue, while spots in cluster 4 are limited to three separate ducts in the right bottom region. The three clusters (cluster 2, 3, and 4) were separated due to difference in their copy number states for regions such as chr2q and chr8q (Supplementary Fig. 11). To illustrate the difference in the copy number profiles between the three clones, the mean fold changes across the spots for each of the three subclones are shown in Fig. 5b. Four regions with significant shifts in the values of at least one of the three subclones were highlighted in the density plots (Fig. 5b; top).

**Figure 5:**
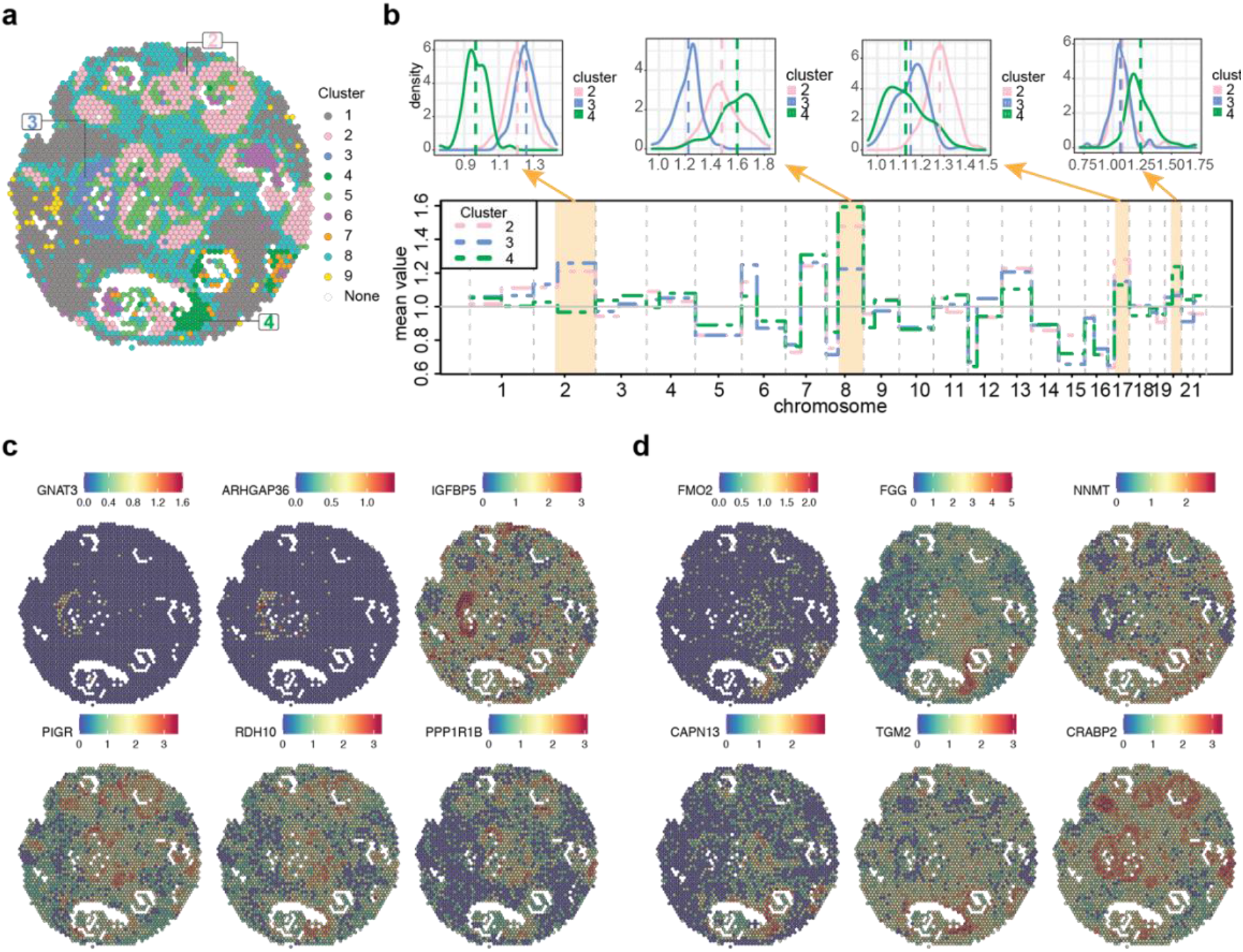
Subclone detection and differential gene analysis on an invasive ductal carcinoma sample. a. Subclones detected from Clonalscope on the ST dataset from an invasive ductal carcinoma sample. b. The mean copy number profiles across the cells assigned to the three spatially segregated subclones (bottom), and the density plots of four regions with significant differences in the copy number states for at least one of the three subclones (top). c. and d. show the top 6 highly expressed genes for cluster 3 and cluster 4. The color scale indicates log-normalized UMI counts for each gene.

Next, we performed differential expression analysis on the three subclones detected by Clonalscope. Gene expression of the top 6 highly expressed genes for subclone 3 and subclone 4 are shown in Fig. 5c and Fig. 5d respectively. Expression of these genes corresponds well to spatial distributions of the two subclones. These genes were also discovered to be associated with outcomes of therapies and different adaptability of tumors. For example, *FGG* is highly expressed in subclone 4. Previous study discovered that breast cancer patients with expression of this gene responded poorly to chemotherapy and had poor survivals ^28^. Another example is the highly expressed *IGFBP5* gene in subclone 3. Expression of this gene is used in the Mammaprint prognostic clinical test to stratify relapse risk for breast cancer patients and to inform therapy decisions ^29^. This gene was also found to be associated with metastatic phenotype in breast tumors ^30^. Higher expression of other genes in the two subclones such as *NNMT*, *TGM2*, and *CAPN13* were all shown to be associated with poor survival and drug resistance in breast cancer patients ^31–33^. Expression of ESR1, PGR, and ERBB2 are routinely tested as clinical biomarkers in breast cancer and their expression did not differ across subclones (Supplementary Fig. 13). The results suggest that the subclones detected by Clonalscope can potentially be the sources of resistance to drug and therapy.

## Discussion

With the development of single-cell sequencing technologies, there is increased interest in subclone detection based on chromosomal copy numbers estimated at the single cell level. Existing copy number-based subclone detection methods are mostly based on hierarchical clustering, and do not leverage information from matched DNA-seq data. Here we presented Clonalscope, a subclone detection method for single-cell and ST data that leverages prior information from matched bulk DNA-seq data for malignant cell/spot labeling and subclone tracing between related datasets. We demonstrated Clonalscope on diverse single-cell data types including scRNA-seq, scATAC-seq, and spatial transcriptomic data. Subclone detection in scRNA-seq and scATAC-seq data were benchmarked with matched bulk and single cell DNA-seq data. In spatial transcriptomic data, expert pathology annotation was used to assess the accuracy of malignant spot labeling, and comparisons between adjacent tissue slices was used to assess the reproducibility of spatial subclone mapping.

Clonalscope estimates copy number states by normalizing to a set of diploid cells/spots at the start. It is important to understand that this does not require the classification of all cells into normal versus malignant status, as we only seek to identify a small set of highly confident normal cells/spots to initialize the normalization. The diploid cells can be a user-selected set of well-known immune or stromal cell types that are easily identifiable with marker genes in standard single-cell data analysis pipelines. Alternatively, Clonalscope can first normalize each cell by the median profile across all cells in the dataset, and then use the malignancy labelling in each round of estimation to iteratively update the diploid cell selection. For ST datasets, diploid spots can be selected based on marker genes, similar to that for scRNA-seq data. Since H&E images are also provided for the ST data, expert pathology annotation can also be useful in verifying/selecting spots from normal regions.

Subclone detection in Clonalscope is based on a nested Chinese Restaurant Process that captures the special, nested structure of subclonal copy number alterations. Within a tumor, CNAs occurring at the early developmental stage are present in most malignant cells, while some subclonal CNAs are only observed in a subset of malignant cells. The nested Chinese Restaurant Process not only samples subclonal membership, but also samples copy number states for each region to create a new copy number profile for the membership assignment. In sampling copy number states, most regions retain the pre-existing state, while there is a rare chance that a region moves to a new state, mimicking the occurrence of a random mutation. The raw outputs from the nested Chinese Restaurant Process depend on user parameters. For more stable estimation, we include a post-processing step that constrains the detected clusters to at least *n_min_* cells.

We also demonstrated the application of Clonalscope to haplotype-informed subclone detection on scATAC-seq data. There is currently a need for reliable haplotype-based subclone detection method for sparse single-cell sequencing data such as scATAC-seq. Improving upon ^9^, Clonalscope does not rely on the known subclones detected from the matched scDNA-seq data. Comparatively, most scRNA-seq protocols only sequence the 3’ or 5’ end of transcripts, limiting allele-specific CNA detection in scRNA-seq data. Recent technologies enable multi-modal sequencing such as RNA and ATAC at the single-cell level ^34^, providing opportunities for broader applications of Clonalscope in analysis of allele-specific CNA profiles.

On the ST dataset from an IDC sample, our detailed analysis identified minor subclones (spots with the same copy number configuration) that might be resistant to therapy. Since this data is not single-cell resolution (each spot contains 1 to 10 cells), Clonalscope also identified clusters consisting of spots with mixtures of tumor and stromal cells based on the gradients of the copy number values. These spots can be the regions of interest for characterizing the interactions between tumor and stromal cells with the help of recently developed cell type deconvolution methods ^35–38^.

Another unique feature of Clonalscope is its ability to trace subclones, a common analysis step in studies where multiple tissue samples are sequenced for the same patient. In this paper, we demonstrated subclone tracing in a data set involving adjacent tissue slices from the same patient. Clonalscope can also be used to link subclones between samples from different time points or different anatomic positions (*unpublished*). Tracing subclones between sequential time points enables the study of temporal dynamics of subclones over the treatment course of cancer, possibly identifying subclones that are resistant to therapy.

## Materials and Methods

### Datasets and pre-processing

The four 10x scRNA-seq datasets, five Visium spatial transcriptomics (ST) datasets, and the SNU601 scATAC-seq dataset analyzed in this study are summarized in Supplementary Table 1. P5847, P6198 and P8823 scRNA-seq data were generated using the protocol described in the previous study ^39^. The Cell Ranger v 3.0 (10x Genomics) was used to generate gene-by-cell matrices of unique molecular identifier (UMI) counts for scRNA-seq data by aligning to the GRCh38 reference genome. The ST datasets of the two replicates from squamous cell carcinoma (SCC) were provided by Ji et al ^23^. The three ST datasets from invasive ductal carcinoma (IDC) were retrieved from the 10x website (https://www.10xgenomics.com/resources/datasets). The SNU601 scATAC-seq dataset was retrieved from our earlier Alleloscope paper ^9^.

### Single-cell and bulk DNA sequencing and data preprocessing

Results of segmentation and subclone detection of the four 10x scRNA-seq datasets were validated with matched scDNA-seq and WGS data (Supplementary Table 3.2). The scDNA-seq data of P5847, P5931 were retrieved from [76] and P6198 were retrieved from [137]. WGS was performed on the P8823 sample for this study.

For WGS sequencing, DNA was first extracted using PureLink™ Genomic DNA Mini Kit as per the manufacturer’s protocols. Genomic DNA was quantified using the Qubit 2.0 Fluorometer (ThermoFisher Scientific, Waltham, MA, USA). For library preparation, NEBNext^®^ Ultra™ DNA Library Prep Kit for Illumina, clustering, and sequencing reagents was used throughout the process following the manufacturer’s recommendations. Briefly, the genomic DNA was fragmented by acoustic shearing with a Covaris S220 instrument. Fragmented DNA was cleaned up and end repaired. Adapters were ligated after adenylation of the 3’ends followed by enrichment by limited cycle PCR. DNA libraries were validated using a High Sensitivity D1000 ScreenTape on the Agilent TapeStation (Agilent Technologies, Palo Alto, CA, USA), and were quantified using Qubit 2.0 Fluorometer. The DNA libraries were also quantified by real time PCR (Applied Biosystems, Carlsbad, CA, USA). Then the sequencing library was clustered onto a flowcell. After clustering, the flowcell was loaded onto the Illumina HiSeq or equivalent instrument according to manufacturer’s instructions. The samples were sequenced using a 2×150bp Paired End (PE) configuration. Image analysis and base calling were conducted by the HiSeq Control Software (HCS). Raw sequence data (.bcl files) generated from the Illumina machine was converted into fastq files and de-multiplexed using Illumina bcl2fastq 2.17 software. One mis-match was allowed for index sequence identification.

### Segmentation on matched bulk DNA-seq data

The first step of Clonalscope is segmenting the genome into regions with different CNA profiles. In this step, matched bulk DNA sequencing data (WGS/WES) or pseudo-bulk data from scDNA-seq data can be segmented by an HMM-based segmentation method. The HMM method operates on the binned counts of pooled cells, and assumes a Markov transition matrix on four hidden states representing deletion, copy-neutral state, single-copy amplification and double-copy amplification: 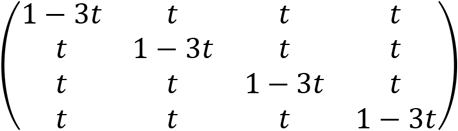, where *t* = 1 × 10^-6^ by default. Emission probabilities follow a normal distribution with 0.2 standard deviation with means depend on individual datasets. The segmentation was performed on pseudo-bulk scDNA-seq data for P5847, P5931, and P6198; WGS data for P8823; and WES for SCC P6. Without matched bulk DNA-seq data, the segmentation was performed on chromosome arms.

### Selection of diploid cells

To identify diploid cells in the scRNA-seq data, the raw scRNA-seq UMI count matrices were first filtered to include cells with at least 500 UMIs, at least 300 genes expressed, and at most 20% of the reads mapped to mitochondria transcriptome. Next, the matrices were normalized with size factors ^40^ and log-transformed. Then PCA was performed, and 30 PCs were selected for clustering and UMAP analysis following the pipeline in the previous study ^41^. For separate clusters on the UMAP, we annotate the cell types in the gastrointestinal tumor tissues using the marker genes from the previous study ^39^. To confidently differentiate malignant cells from normal epithelial cells in the P5931 scRNA-seq dataset, we processed both tumor and normal scRNA-seq datasets together using the same pipeline described above, and successfully labeled normal epithelial cells in the tumor dataset in a separate cluster mainly containing cells in the normal dataset (Supplementary Fig. 4).

To identify reference cells in the Visium ST datasets, we first ran the Seurat (4.0.1) ^42^ pipeline like that for scRNA-seq data to retrieve some preliminary clusters at the spot level. Since the resolution of the Visium datasets are 1 −10 cells, marker gene expression is not locally enriched but gradient. However, we could still select a cluster with least expression of the genes in cancer cells (eg. ERBB2 for breast cancer cells) and high expression of other marker genes for diploid cells. With the help of H&E images, we could also more confidently select the clusters located in the regions mainly consisting of diploid cells.

### Computing coverage change

With each region *r* identified from matched DNA-seq data, we applied a Poisson model to reduce a gene-by-cell matrix to a region-by-cell matrix for scRNA-seq data. For cell *i* and gene *g*, we defined the raw UMI count matrix as *N_ig_* with

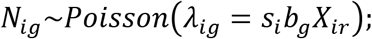

where *s_i_* is the normalizing factor of cell *i*; *b_g_* is the baseline expression of gene *g*; and *X_ir_* is the relative fold change of region *r* for cell *i*. The normalizing factor *s_i_* is computed as follows:

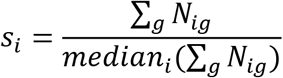

With a set of control cells being defined, *δ_ir_’s* for these cells are equal to 1. Therefore, *b_g_* can be estimated by solving

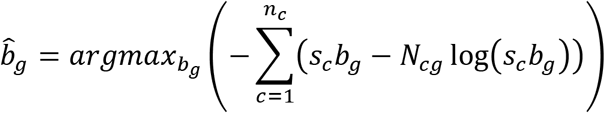

By derivation, 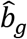 can be solved with direct computation.

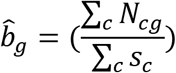

Then for each cell *i*. and region *r*,

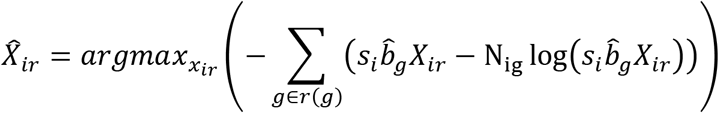

where *r*(*g*) is region *r* which gene *g* belong to.

By derivation, 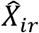 can be solved with direct computation.

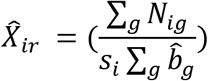

For ST datasets, the model is the same, but *i* is for spot instead of cell.

In the real estimation, we excluded the cycle genes, HLA genes and the mitochondria genes learning from CopyKAT ^15^. We also filtered out the extremely variable genes and regions overlapping <150 genes.

### Subclone detection with Nested Chinese Restaurant Process

After the relative fold changes for each cell and each region 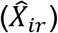 are retrieved, A Nested Chinese Restaurant Process was applied to detect subclones in scRNA-seq data in a non-parametric manner. First, we consider the coverage-only mode. The model for *X_ir_* is

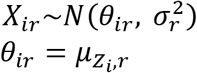

with *z_i_* ∈ {1,…, *K*] is subclone *k* which cell *i* is assigned to. *μ_kr_* is the relative fold change of subclone *k* in region *r*, and *σ_r_* is the standard deviation of region *r*. The targeted outputs of the Nested Chinese Restaurant Process are *Z_i_*, the estimated subclonal identity of cell *i*; and *U_kr_*, the copy number states of region *r* for subclone *k*.

To include the prior information from the matched bulk DNA-seq data, we first set the prior *U_prior_* as two clusters

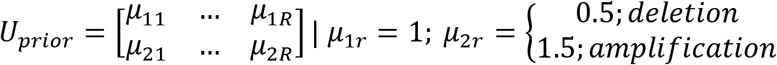

with *μ_2r_* known from the DNA-seq data. Then for each cell *i* = 1~*N*,

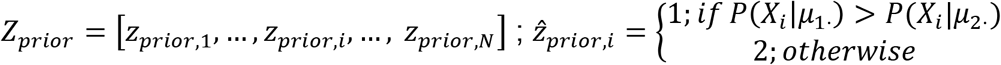

In the *t^th^* iteration of the Nested Chinese Restaurant Process, we first iterate over all cell *i*. To assign cell *i*. into one of the existing clusters, the probability of

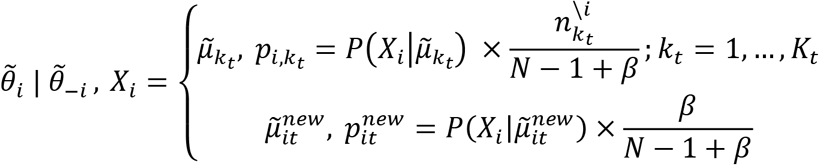

where 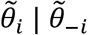, *X_i_* is the variable for the estimated mean of relative fold change for cell *i*. across all the regions, conditioning on all other cells and the observation. 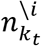 is the number of cells assigned to cluster *k* from the *t^th^* iteration excluding cell *i*; and *N* is the total number of cells. *β* is the scalar parameter controlling the chance of accepting a new cluster in the process. *β* was by default set as 2. 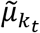 is the relative fold changes across all regions for subclone *k* form the *t^th^* iteration, and *p_i,k_t__* is the probability of cell *i* being assigned to the *k_t_* cluster. 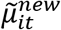 is the relative fold changes across all regions for a newly seeded cluster for cell *i* in iteration *t* with the probability being 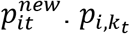 and 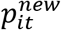 can be calculated as followsy

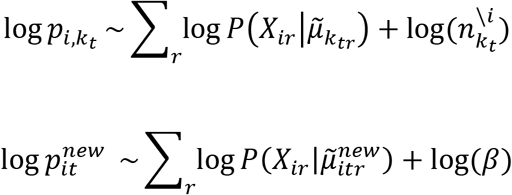

To create 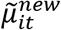, the second Chinese Restaurant Process was executed to select one of the existing copy number states or seed a new copy number state for each region. We skip the subscript “it” for the following notation. For each region *r*,

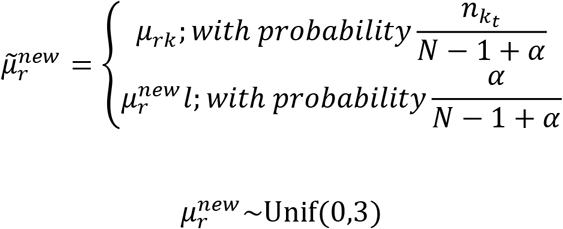

where 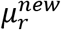 is a newly seeded copy number state for region *r*. *n_k_t__* is the number of cells assigned to cluster *k* from the *t^th^* iteration. *α* is the scalar parameter controlling the chance of accepting a new cluster in the process. By default, *α* is set as 2. With 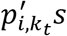 for all *k_t_* clusters and 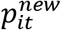 being calculated, 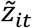 is sampled based on the probability.

After 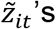 are updated for all cell *i*, *μ_kr_* is sampled based on the following distribution

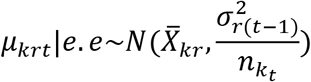

where *σ_r(t–1)_* is the standard deviation of region *r* for the (*t* – 1)^th^ iteration. Then *σ_rt_* can be updated by computing

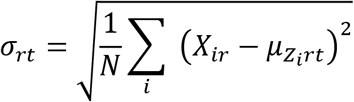

For subclone detection with both coverage and allelic ratio, the model for *X_ir_* is

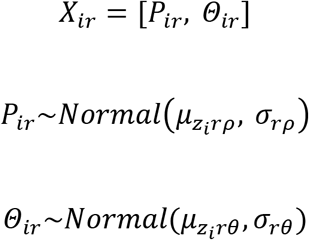

In the formula, *P_ir_* and *Θ_ir_* are the estimated coverage and major haplotype proportion ^9^ for region *r* in cell *i*. *z_i_* ∈ {1,…, *K*] is subclone *k* which cell *i*. is assigned to, and *μ_krρ_* and *μ_krθ_* are the relative fold change and major haplotype proportion of subclone *k* in region *r* respectively. *σ_rρ_* and *σ_rθ_* are the standard deviations of relative fold change and major haplotype proportion for region *r*. The Nested Chinese Restaurant Process is similar to that for the coverage-only mode. The outputs from the estimation are *Z_i_*, the estimated subclonal identity of cell *i*; and [*U_krρ_, U_krθ_*] the allele-specific copy number states of subclone *k* for region *r*.

The two parameters *α* and *β* control the numbers of subclones detected from the Nested Chinese Restaurant Process. To retrieve more stable estimation, the first few iterations of the raw outputs from the Nested Chinese Restaurant Process are trimmed. By default, 200 iterations are run and the estimates from the first 100 iterations are trimmed. After the trimming step, each cell is assigned into the most possible subclone *k* using the majority vote strategy. Next, only the subclones with > *n_min_* cells assigned are kept, and the smaller subclones are removed. By default, *n_min_* is set as 1 % of cells. Then the cells originally assigned to the removed subclones are re-assigned to the existing subclones. Then *U_kr_* the copy number states of region *r* for subclone *k* is updated based on the re-assignment.

### Assessing segmentation on scRNA-seq data and on matched DNA-seq data

To evaluate different segmentation methods on scRNA-seq data, we calculated the correlations between the mean copy number profiles in scRNA-seq data to those from (pseudo)bulk DNA-seq data. Three segmentation methods for scRNA-seq data were compared—segmentation from matched bulk DNA-seq data; segmentation from CopyKAT and segmentation from inferCNV. CopyKAT and inferCNV were ran using the default parameters. CopyKAT and inferCNV perform segmentation mainly at the gene level, and they output the estimated copy number profiles for each gene in each cell. Therefore, we computed the mean copy number profiles across all epithelial cells for each gene. Since the output values for CopyKAT are at the log-scale, the values were exponentially transformed to original values. Comparatively, we used Clonalscope to perform segmentation on matched DNA-seq data and compute the mean fold changes for each segment in scRNA-seq data. For easier comparison with CopyKAT and inferCNV, the mean fold changes for each segment were transformed to the gene level.

All scDNA-seq data were pre-processed with Cell Ranger DNA pipeline (https://support.10xgenomics.com/single-cell-dna/software/overview/welcome, beta v.6002.16.0). For P5847 and P5931, we applied Alleloscope ^9^ on the scDNA-seq data, and computed the mean coverage changes across all identified malignant cells for each segment to retrieve the mean copy number profiles for the pseudo-bulk DNA-seq data. For P6198, we generated the pseudo-bulk sample from the scDNA-seq data, and computed the mean coverage changes for each segment. For P8823, we first quantified the reads in each window with a 100000 bp size for both matched tumor and normal WGS data by running the “intersect” command in the bedtools ^43^. The profile of the mean coverage changes for P8823 bulk DNA-seq data were then computed for each segment. Similarly, we also transformed the vector of the mean coverage changes for all DNA-seq data from the segment scale to the gene scale.

### Malignant cell/spot labeling on scRNA-seq and spatial transcriptomic data

Copy number profiles from matched bulk DNA-seq data can also be used to reliably detect malignant cells/spots in scRNA-seq and ST datasets. After the subclones are detected in the scRNA-seq and ST datasets, 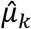, the estimated copy number profiles of subclone *k* can be compared to the copy number profiles from matched DNA-seq data (*μ*_1_) across all regions. To determine subclone *k* is malignant or not, the revised cosine similarity (*S_c_*) was computed as

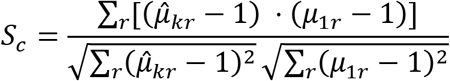

If *S_c_*>*S_min_* (0.5 by default), the subclone *k* is labeled as malignant.

Accuracy of malignant cell labeling was compared between Clonalscope, CopyKAT, and hierarchical clustering on an scRNA-seq dataset. Clonalscope and CopyKAT were run using default parameters. For hierarchical clustering, we first retrieved segments from matched DNA-seq data, and estimated relative coverage fold changes for each cell in each segment using the function of Clonalscope. Then hierarchical clustering was applied to cluster the cells based on the estimated copy number profiles versus the Bayesian non-parametric clustering used in Clonalscope.

### Comparison of subclones between scRNA-seq data and matched scDNA-seq data

On the malignant cells identified in the P5931 scRNA-seq data, we compared the subclones detected by Clonalscope, inferCNV, and copyKAT to those in the matched scDNA-seq data. In the scDNA-seq data of P5931, 3 major subclones (D1~D3) were observed, which can be used as the ground truth of the sample. For P5931, since the matched pseudo-bulk DNA-seq data suggested that CNAs were mainly located on four chromosomes—chr7,8,20,21, Clonalscope was run on these four chromosomes to detect subclones with default parameters. For hierarchical clustering used in CopyKAT and inferCNV, the numbers of subclones need to be specified. Since 6 and 5 subclones were used in the segmentation for CopyKAT and inferCNV, we set the output numbers of subclones as 6 and 5 respectively.

After the subclones in scRNA-seq data were detected from the three methods, we mapped the subclones to the corresponding “true” subclones observed in the matched scDNA-seq data. For subclone mapping, the revised cosine similarity (*S_c_*) was computed:

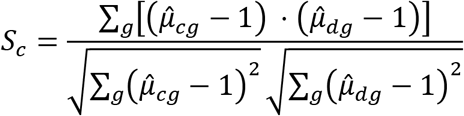

where 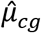 and 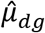 are the mean fold change for gene *g* across the cells assigned to subclone *c* in scRNA-seq data and subclone *d* ∈ [*D*1, *D*2, *D*3] in scDNA-seq data. Subclone *c* in scRNA-seq data was mapped to the subclone *d* with the highest similarity score. For Clonalscope, we summed over all the region *r* instead of gene *g*.

### Subclone detection in scATAC-seq data

Following the pipeline described in our previous study^9^, we processed the SNU601 scATAC-seq data and generated the region-by-cell matrices for relative fold changes and allelic ratios. Next, we constrained the values for the relative fold changes to be < 3, and the values for the allelic ratios > 0.05 and < 0.95 for more stable estimation. In our previous study, six major subclones were detected in the SNU601 sample with difference in their allele-specific copy number profiles mainly on 10 regions. Therefore, we selected the 10 regions for the downstream subclone detection. Cells with no reads detected in more than five out of the ten regions were filtered out. With the two processed matrices as the input, Clonalscope was ran to detect the subclones based on both coverage changes and allelic ratios using the default parameters. The final clusters were requested to include > 50 cells. The subclones detected from Clonalscope were compared to the supervised clustering used in the previous study, which requires a known number of subclones and their corresponding copy number profiles from scDNA-seq data. The two methods were shown parallelly for easier comparison.

### WES data processing

The bam file of the WES data from SCC P6 was retrieved from GSE144240. On the bam files for matched tumor and normal samples, we first quantified the reads in each window with a 100,000 bp size by running the “intersect” command in the bedtools ^43^. Then we segmented the genome into regions with the same copy number states using the HMM segmentation function in Clonalscope. Since the dataset was originally aligned to hg19, we used the LiftOver tool (https://genome.ucsc.edu/cgi-bin/hgLiftOver) to convert the segments to GRCh38 to be compatible with the matched ST dataset.

### Spatial transcriptomic (ST) data analysis

Clonalscope was run to detect subclones without considering spatial loci in ST datasets. The estimated clonal identity of each spot can then be visualized spatially, enabling integration of the two types of signals. Seurat was used to align each spot to the corresponding loci in the H&E image.

For SCC P6 sample, the copy number profiles of the identified malignant spots in the two replicates were compared to each other by computing the correlation. The spatial distributions of the malignant spots were also shown. On the IDC sample with 2 sections, to trace subclones from one ST dataset to another ST dataset, we used the subclones detected on the first section (section 2) as the prior clusters for Clonalscope’s subclone detection on another section (section 1). The prior clusters can be easily provided to Clonalscope, and the direct information from bulk DNA-seq data is ignored in the mode. On the IDC sample with three subclones detected (Fig. 5), we assessed the difference in the mean copy number profiles of the three clones. Differential gene expression analysis on the three clones was performed using the “FindMarkers” function in Seurat ^42^. Each of the three subclones were compared against the other two subclones. The expression values of each feature were visualized using Seurat.

## Data availability

The patient scRNA-seq and WGS data generated for this study are available under dbGAP identifier phs001818. Data is available with dbGaP authorized access for Health/Medical/Biomedical purposes. The other patient scRNA-seq data were obtained from dbGAP under accession phs001818 ^39, 44^. The patient scDNA-seq data were from dbGAP under accession phs001711 ^9^ and phs001818 ^44^. For the SNU601 sample, scDNA-seq data and scATAC-seq data were retrieved from the SRA under accession PRJNA598203 ^45^ and PRJNA674903 ^9^ respectively.

## Code availability

Clonalscope is available on GitHub at https://github.com/seasoncloud/Clonalscope.

## Acknowledgements

The work is supported by the National Institutes of Health [P01HG00205ESH to B.T.L., S.M.G. AND H.P.J., 5R01-HG006137-07 and 1U2CCA233285-01 to C-Y.W. and to N.R.Z.]. Additional support to HPJ came from the Research Scholar Grant, RSG-13-297-01-TBG from the American Cancer Society, Clayville Foundation and the Gastric Cancer Foundation.

## Author contributions

C.-Y.W. and N.R.Z. conceived the computational methods and designed the study with help from H.P.J. C.-Y.W. developed and implemented the computational methods and conducted all data analyses. A.S. performed all related sample preparation and sequencing. J.R. helped with benchmarking and improvement of the method. P.R.H. helped with pathology annotation and data interpretation. B.T.L. helped in data interpretation. S.M.G. performed data pre-processing and coordinated data transfer. H.P.J. advised all experiments and data collection. C.-Y.W. and N.R.Z wrote the manuscript. All authors read and approved the final manuscript.

