## Supplementary file for "Cancer subclone detection based on DNA copy number in single cell and spatial omic sequencing data"

### Supplementary Information

### Supplementary Figures

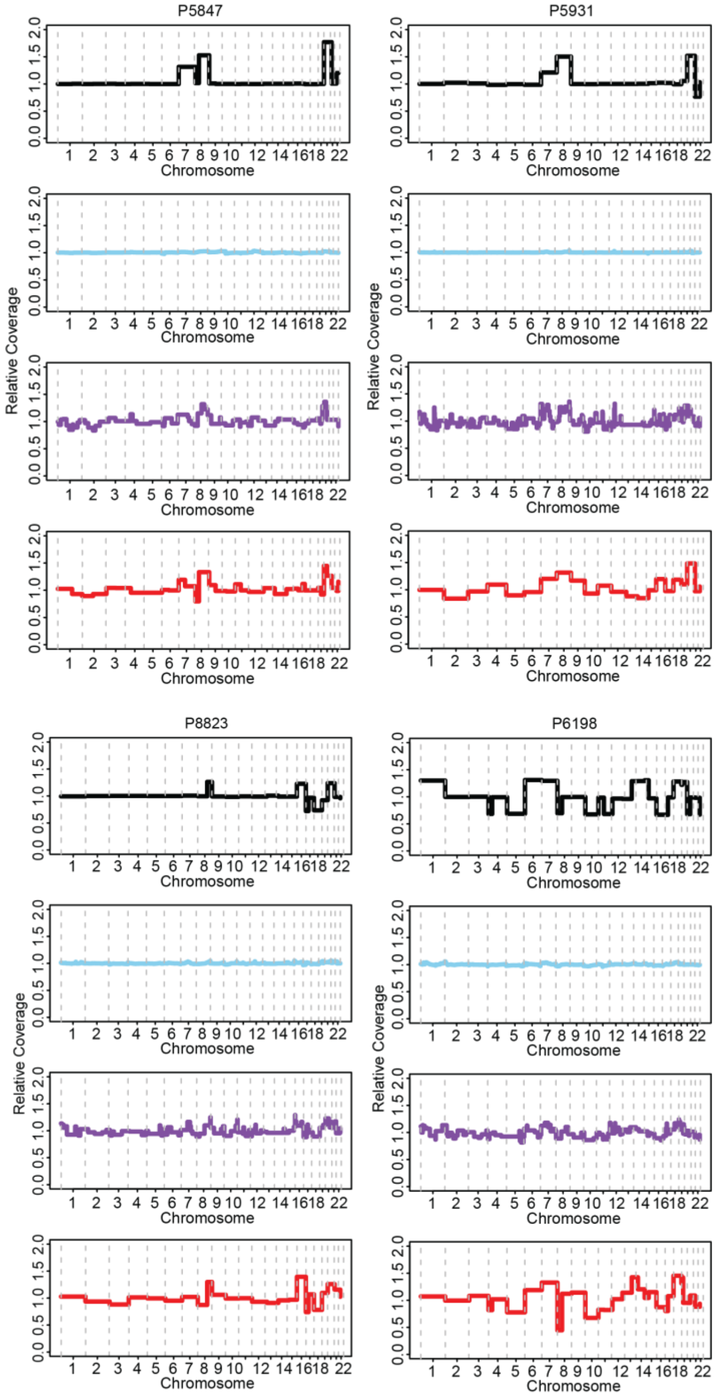

***Supplementary Figure 1: Comparison of the mean coverage changes using segments from different methods to those observed in matched (pseudo)bulk DNA-seq data on four gastrointestinal samples.***

The mean coverage changes in scRNA-seq data using segments from inferCNV (sky-blue), CopyKAT (purple), and bulk DNA-seq data (red) were compared against the mean coverage changes in DNA-seq data (black). Four segmentation plots are aligned for each sample. The four samples are P5847 (top left), P5931 (top right), P8823 (bottom left), and P6198 (bottom right).

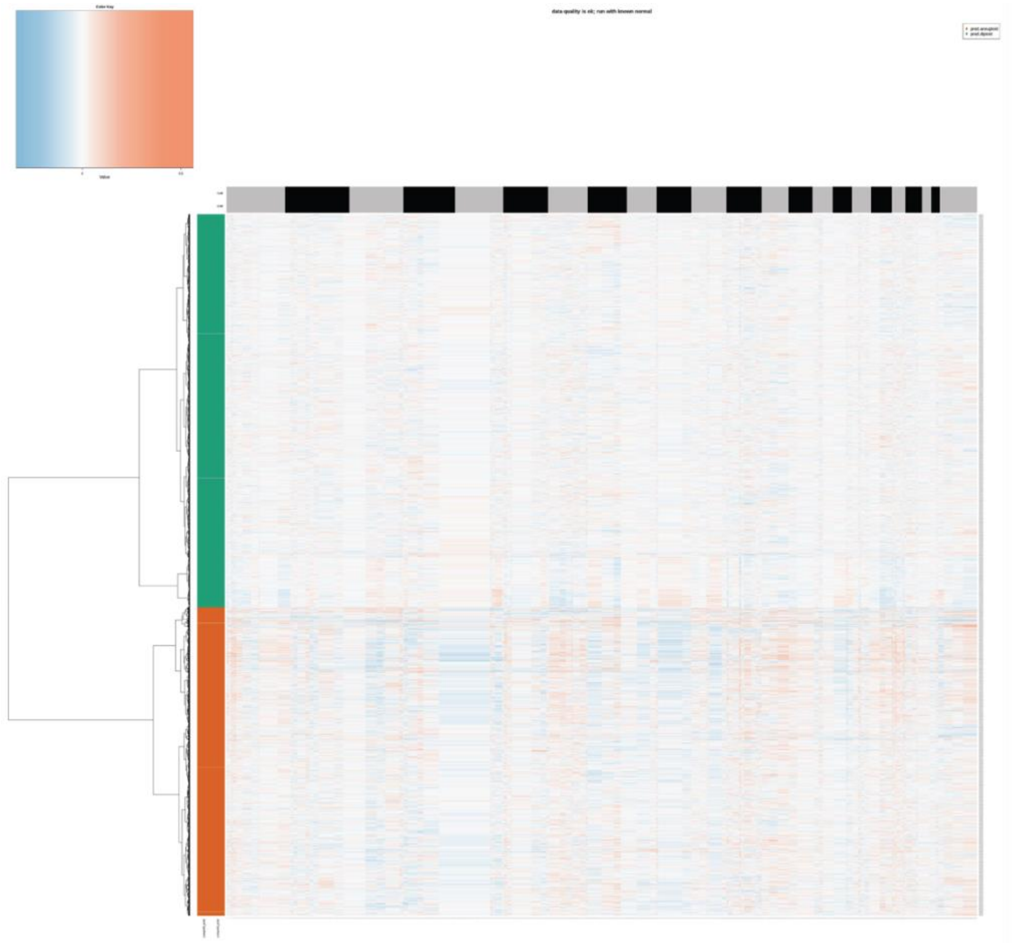

***Supplementary Figure 2: Malignant cell labeling result from CopyKAT for the P6198 scRNA-seq data.***

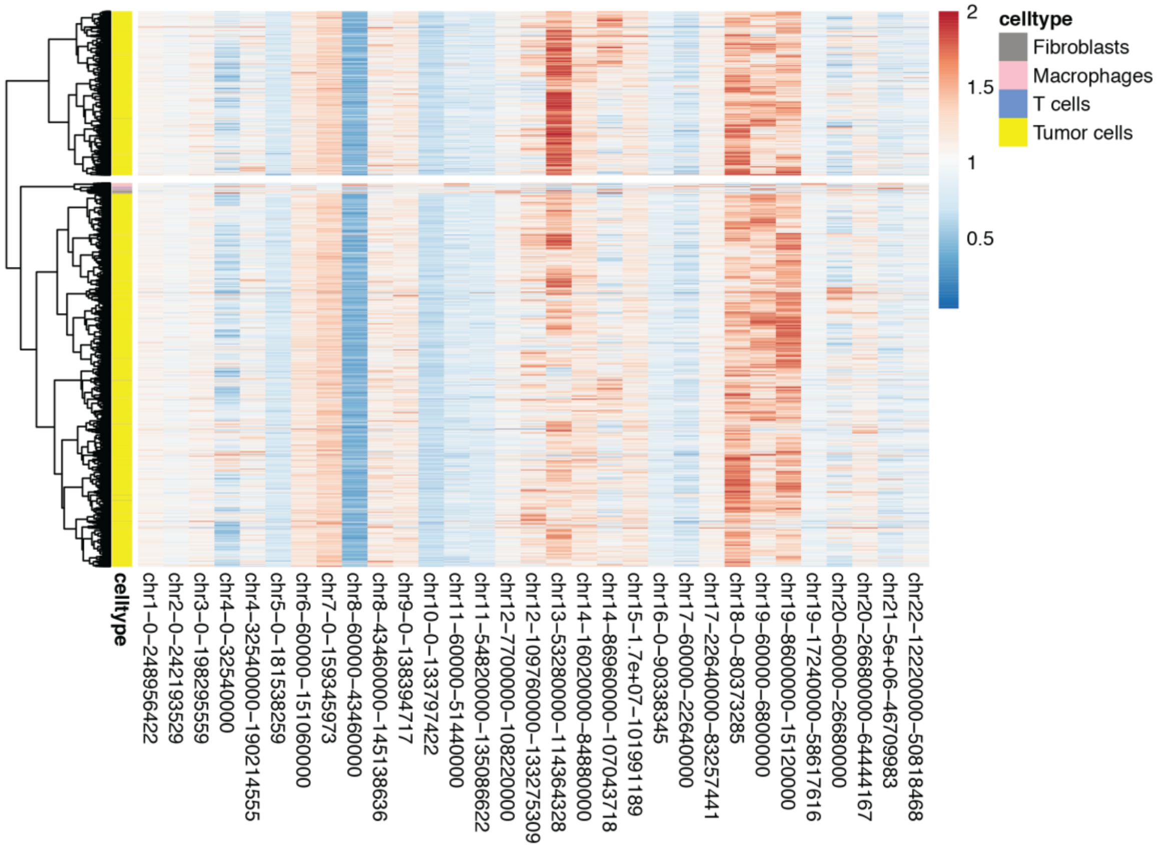

***Supplementary Figure 3: Hierarchical clustering result on the values of fold changes for each cell and each segment for the P6198 scRNA-seq data.***

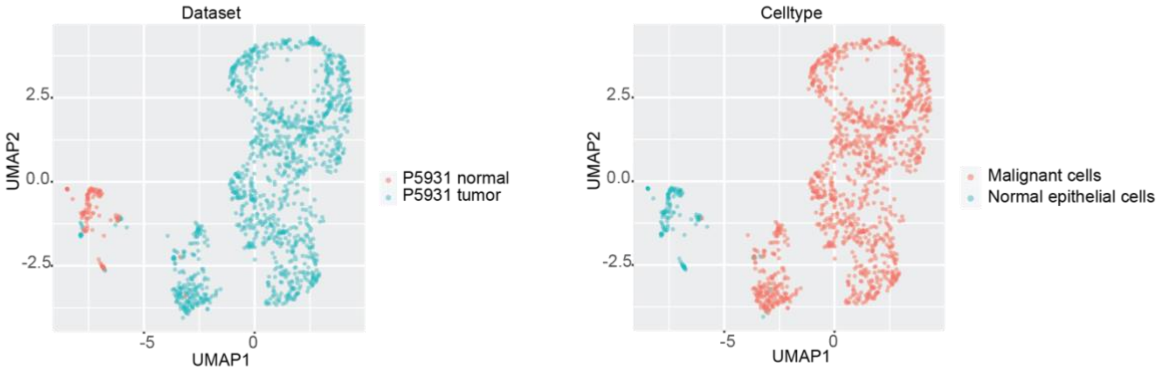

***Supplementary Figure 4: UMAP plots for the epithelial cells in matched tumor and normal scRNA-seq data from the P5931 sample.***

The matched tumor and normal scRNA-seq data were analyzed together for dimensionality reduction, clustering and cell-type annotation.

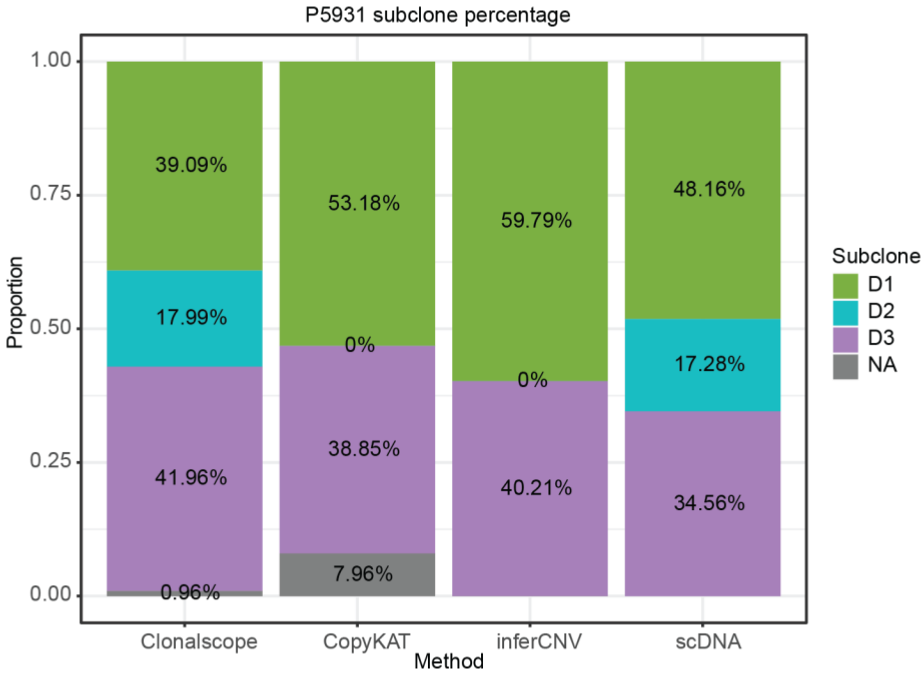

***Supplementary Figure 5: Proportions of the subclones detected from Clonalscope, CopyKAT, and inferCNV compared to those observed in the matched scDNA-seq data for the P5931 tumor sample.***

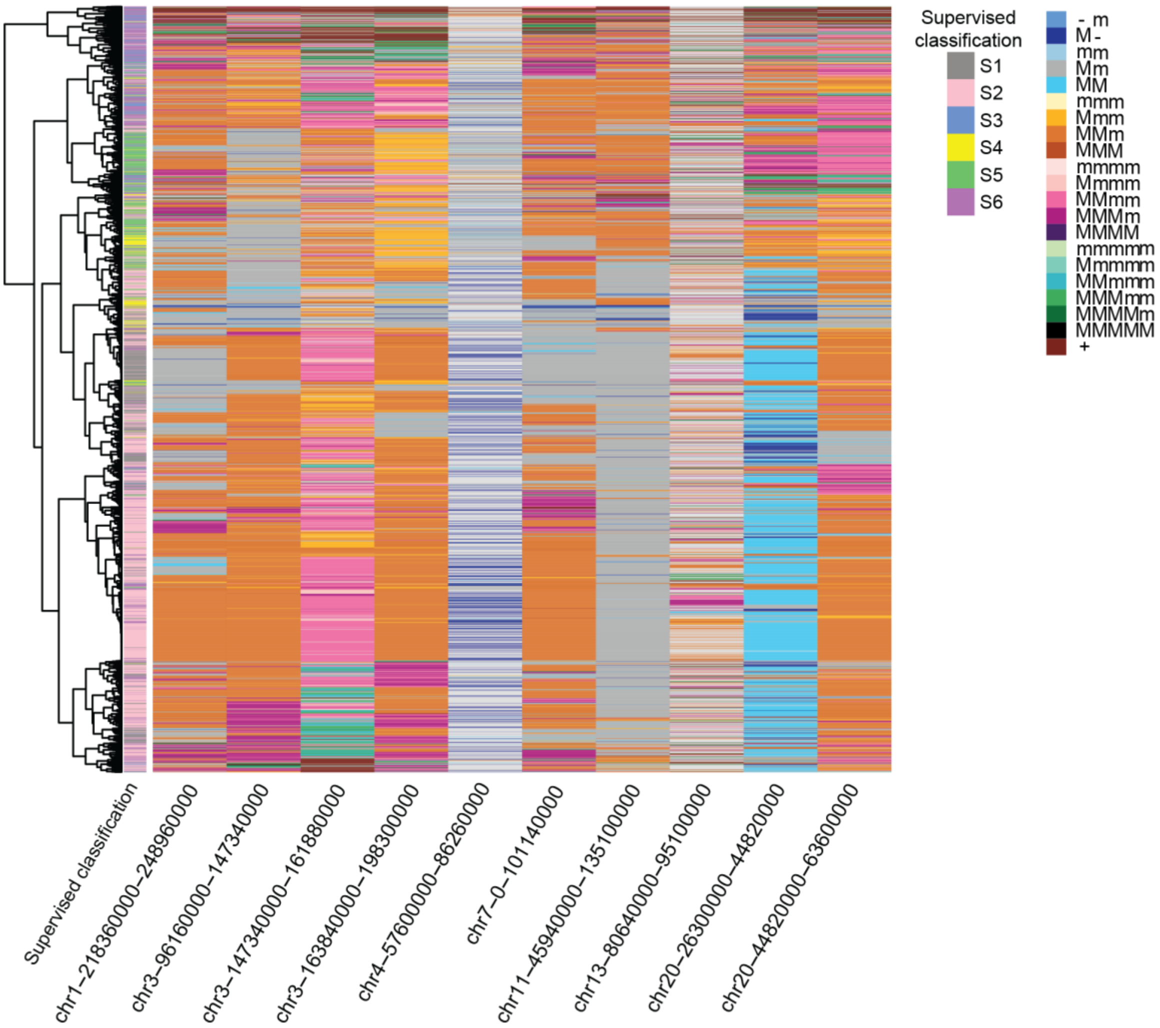

***Supplementary Figure 6: Hierarchical clustering result on the values of fold changes in ten marker regions for each cell in the SNU601 scATAC-seq data.***

The supervised classification is shown in the left. In the color legend, M and m represent the major haplotype and minor haplotype, respectively.

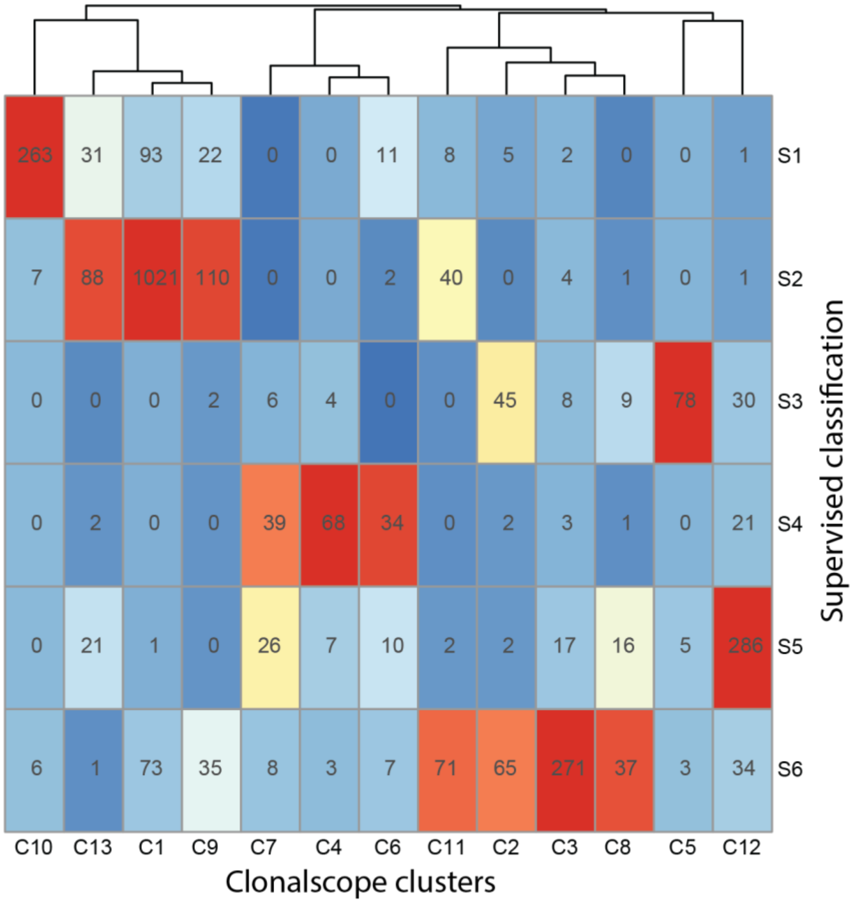

***Supplementary Figure 7: Heatmap showing frequencies for each combination of clusters from Clonalscope estimation and supervised classification.***

The color scale represents the standardized values for each column.

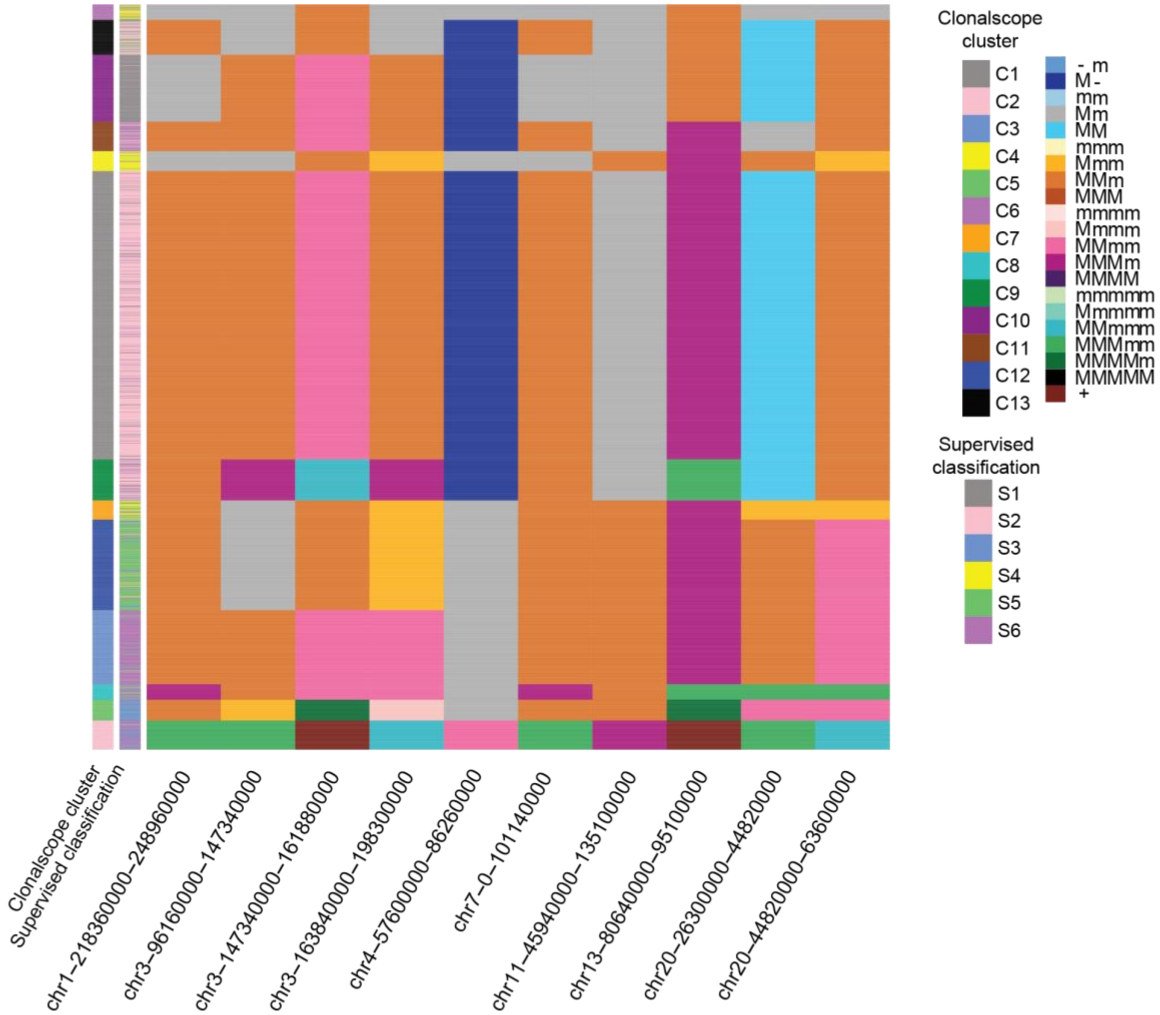

***Supplementary Figure 8: Consensus plot of the subclone detection result from Clonalscope on the SNU601 scATAC-seq dataset.***

The clusters from Clonalscope and supervised classification are shown parallelly in the left. In the color legend, M and m represent the major haplotype and minor haplotype, respectively.

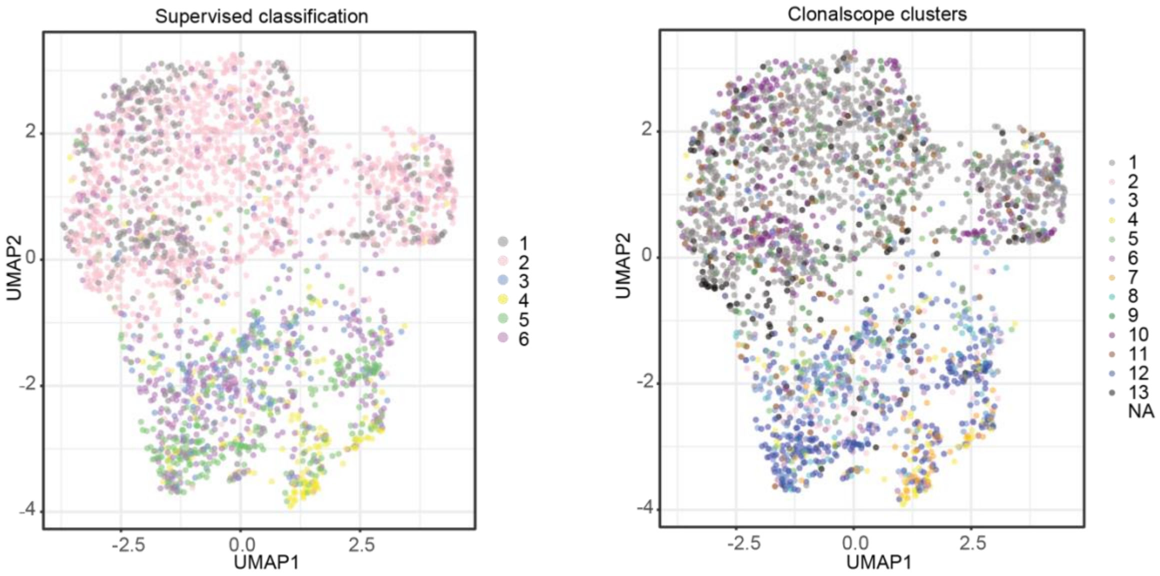

***Supplementary Figure 9: UMAP projection of genome-wide scATAC-seq peak profile on 2,753 cells colored by the clusters from supervised classification (left) and Clonalscope estimation (right).***

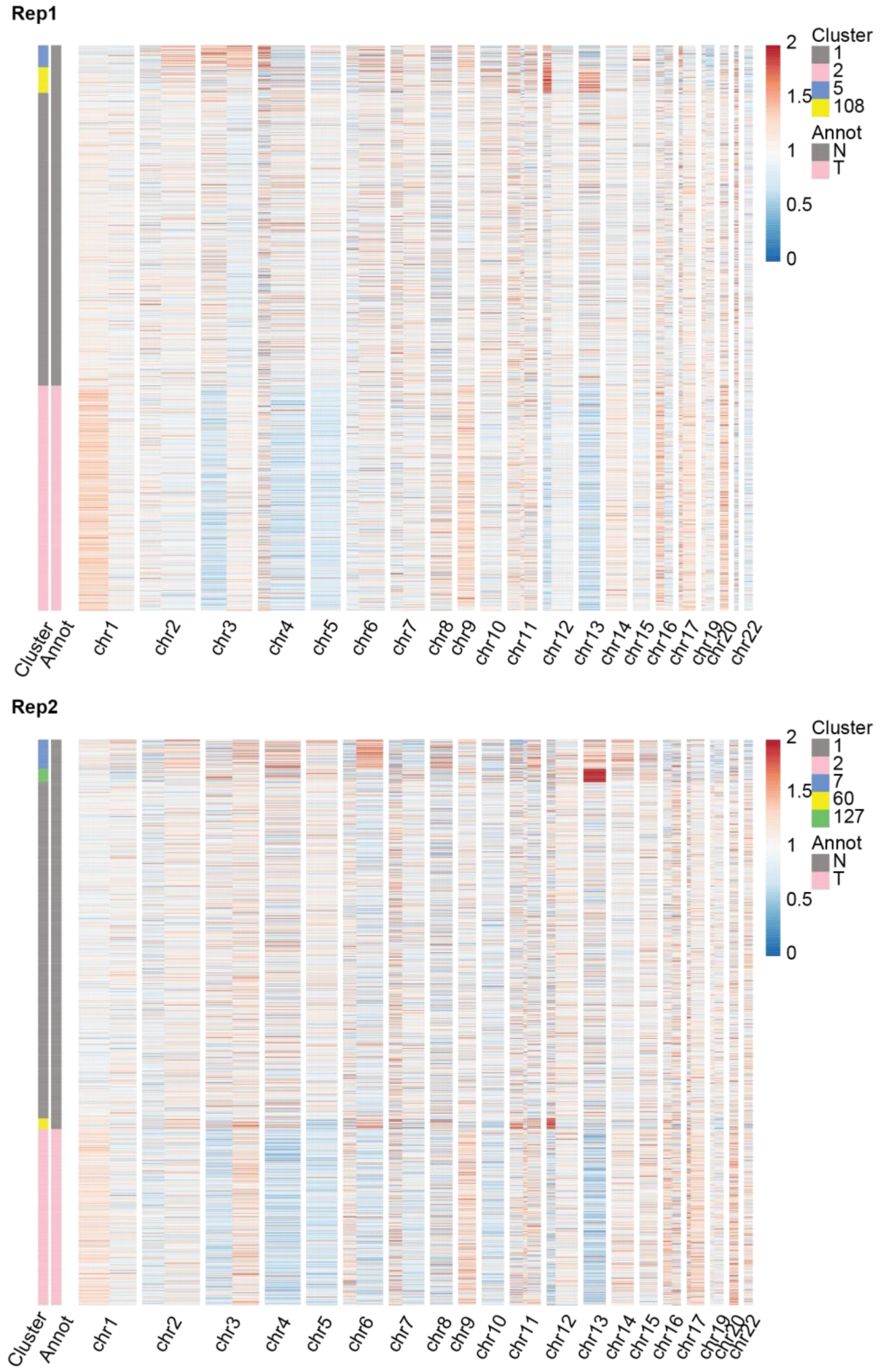

***Supplementary Figure 10: Supplementary Figure 3.10: Heatmaps showing the subclone detection and malignant cell labeling results from Clonalscope on the ST datasets from two replicates of the SCC P6 sample.***

The heatmaps for replicate 1 and replicate 2 are shown in the top and bottom respectively. In the legend, ‘Annot’ represents the annotation of tumor (T) and normal (N) cells.

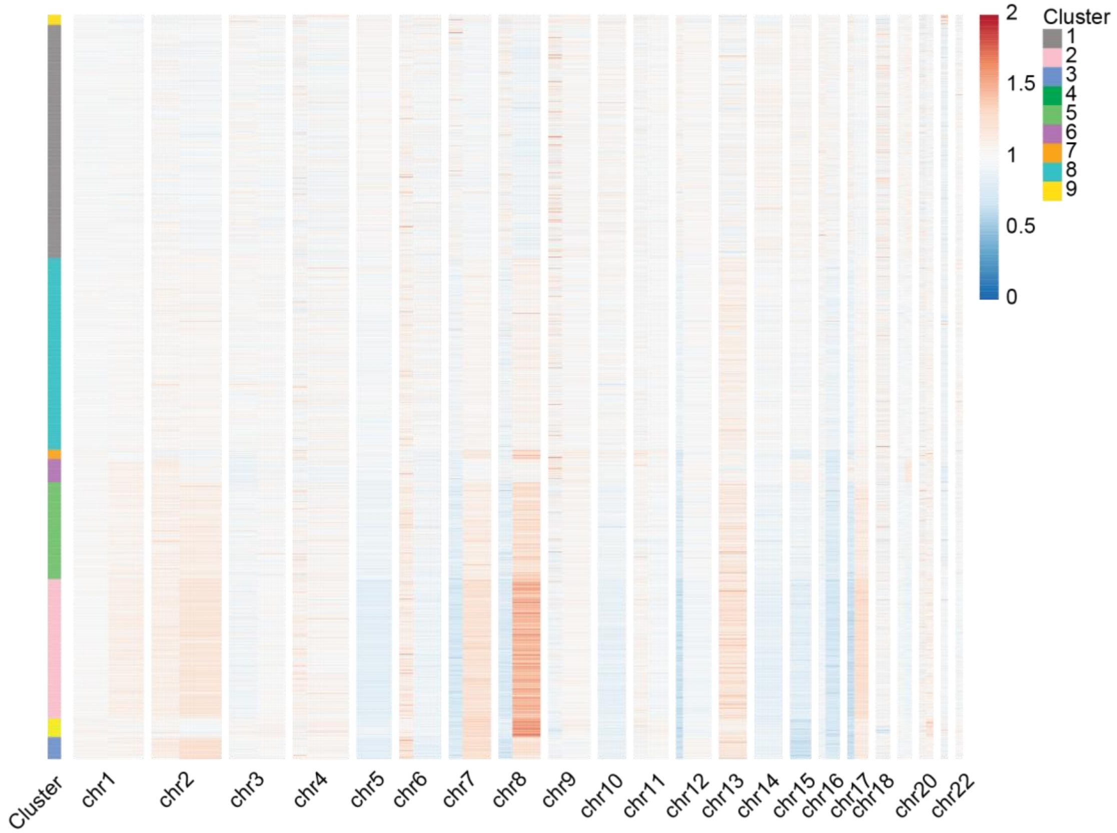

***Supplementary Figure 11: Heatmap showing the subclone detection result from Clonalscope on the ST datasets from an IDC sample.***

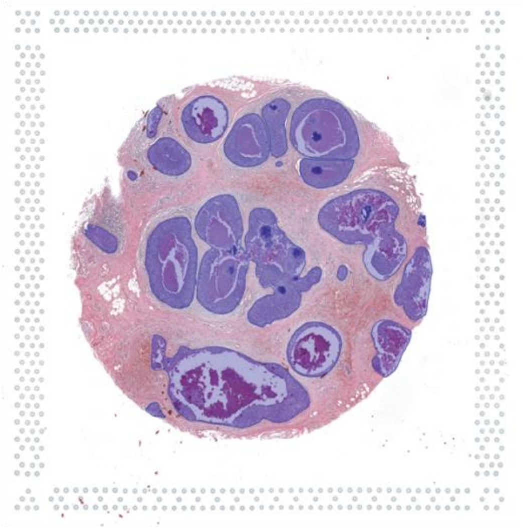

***Supplementary Figure 12: The H&E image of the invasive ductal carcinoma sample with pathology annotation.***

The regions containing malignant cells are annotated with blue overlays.

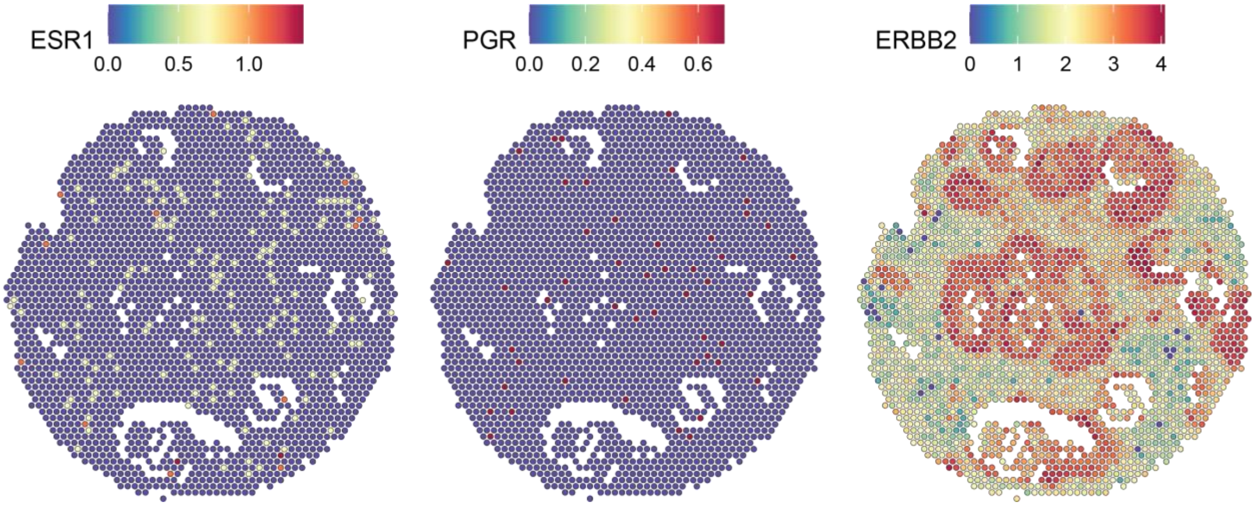

***Supplementary Figure 13: Expression of ESR1, PGR, and ERBB2 genes in each spot.***

The color scale indicates log-normalized UMI counts for each gene.

### Supplementary Tables

***Supplementary Table 1: Summaries of the scRNA-seq and scATAC-seq datasets.***

| Sample | Cancer type | Source | Paired normal | Type | Mean reads per cell | Cell number | Ref |
|---|---|---|---|---|---|---|---|
| P5847 | Gastric | Primary tissue | No | scRNA-seq | 16,184 | 7,387 | This study |
| P5931 | Gastric | Primary tissue | Yes | scRNA-seq | 19,129 | 5,861 | [1] |
| P6198 | Colorectal | Liver meta | No | scRNA-seq | 51,399 | 7,190 | [2] |
| P8823 | Colorectal | Primary tissue | No | scRNA-seq | 26,366 | 26,523 | This study |
| Sample | Cancer type | Source | Paired normal | Type | Median frags per cell | Cell number | Ref |
| SNU601 | Gastric | Ascites meta | No | scATAC-seq | 69,455 | 3,614 | [3] |

***Supplementary Table 2: Summaries of the DNA-seq datasets.***

| Sample | Cancer type | Source | Type of DNA-seq | Coverage per cell | Cell number | Ref |
|---|---|---|---|---|---|---|
| P5847 | Gastric | Primary tissue | scDNA-seq | 422,134 | 715 | [3] |
| P5931 | Gastric | Primary tissue | scDNA-seq | 730,932 | 796 | [3] |
| P6198 | Colorectal | Liver meta | Pseudo bulk | - | - | [4] |
| P8823 | Colorectal | Primary tissue | WGS | - | - | This study |
| SNU601 | Gastric | Ascites meta | scDNA-seq | 565,648 | 1,531 | [5] |
